# Growth bistability in small bacterial populations exposed to antibiotics

**DOI:** 10.64898/2026.05.21.726888

**Authors:** B. Ledoux, D. Lacoste

## Abstract

With the development of microfluidics, it has now become possible to assess the susceptibility of bacteria to antibiotics at the single-cell level instead of relying on population measurements. Such studies are particularly relevant when the growth of bacterial population in the presence of antibiotics is heterogeneous. Here, we build a model to describe such a case, and apply it to experimental measurements on a small population of *E. Coli* exposed to ciprofloxacin, a drug which is well known for triggering a bistable response.

---

The phenomenon of drug-resistance, and in particular antibiotic resistance, represents a challenge to understand because it involves the coupling of the antibiotic drug with the complex metabolism of a cell. Given the complexity of this interaction, simplified toy models of the cell metabolism, even at the single-cell level, are useful and can yield quantitative predictions on the average reduction of bacterial growth by antibiotics [1–3].

One interesting result from this line of work is the prediction of a bistable response in a certain range of antibiotic concentration and for a specific type of antibiotics that binds irreversibly to key molecules of metabolism. This heterogeneous response has been observed in a number experiments [4–7]. In practice, the cell population contains two phenotypically distinct subpopulations [8– 19], corresponding to growing and non-growing or dormant cells, which respond differently to the antibiotics. The drug ciprofloxacin fits with this picture, because it is known to elicit two heterogeneous subpopulations via the SOS response, which is triggered by DNA damage [11, 16, 18–21].

Besides dormancy, antibiotics can also induce the phenomenon of persistence, characterized by the ability of a small fraction of bacterial cells to survive a lethal dose of antibiotics. While it is generally assumed that persistence is caused by growth-arrested dormant cells generated prior to drug exposure, there are also forms of persistence that arise from cells that were growing actively before the exposure to antibiotics [22]. Thus, it appears that there are many different survival pathways for bacteria to respond to antibiotics.

It is nowadays possible to explore these questions using single-cell time lapse experiments and mother machine experiments. In addition to these setups, a novel experimental microfluidic platform has been built [23, 24] to measure the susceptibility of bacteria to antibiotics in small populations. In this device, a large number (*N*_*d*_ > 10^2^) of identical droplets containing nutrients and antibiotics at a controlled concentration *c* are anchored on a plate. Each droplet is then seeded with a small number of bacteria (*E. Coli*), an inoculation step which can be modeled by a Poisson distribution [25, 26], with a controlled mean, which we will take of order 1 here. This form of distribution is justified when all the droplets have the same volume, which is small compared to the total volume of the inoculated solution. Then, droplets evolve independently for a fixed duration *t*, and the number of cells *n* in each droplet is monitored using image analysis pipelines. Thanks to the large number of independent droplets present in the device, there is enough data to measure the statistics of a single bacteria to form small colonies and estimate the probability distribution of the cell population as a function of time and for different antibiotic concentrations.

The experiment shows that this probability distribution is typically a decreasing function of the population *n* for small enough populations, even at times which are long compared to the typical division time [23, 24]. A striking feature of this distribution is the presence of a tail which extends to small populations of about 5 to 6 cells. In a first guess, one could propose that the tail arises from the transient evolution of a birth/death process with fixed birth/death rates. However, the state *n* = 0 is an absorbing state, and no current is coming out this state, which means that one would eventually obtain a finite peak at *n* = 0, with no significant tail for small finite values of *n*.

Another idea would be to assume that the experiment fails to distinguish between bacterial death (the bacteria are no longer observed) from non-growing bacteria. In that case [27], the total number of cells (either dead or alive) is measured over the whole trajectory instead of the number of living bacteria. Thus, there is a subpopulation of individuals—dead bacteria—whose growth and death rates are zero, essentially a dormant state. Indeed, dead cells do not disappear since they continue to be counted (zero death rate) and they are unable to divide (zero growth rate). The transition from the active state to the dormant (or dead) state is irreversible, and there is no flux from non-zero populations to the empty droplet state (see Suppl. Mat. for details). Because dead cells are still counted, the population within a droplet can only increase or stay constant. As a result, the probability to have an empty droplet should remain constant in time (cells cannot appear ex nihilo), which is in contradiction with the observed data.

We now come to the interpretation which we favor, namely that the tail in the cell population results from growth heterogeneity of the population. Instead of considering that the bacterial population is characterized by a single growth rate, we assume that it is composed of discrete phenotypes, characterized by their growth and death rates. Both explanations can be modeled as a population of cells transitioning from one state to another, defined by their growth and death rates, but the second explanation is better supported by experimental data, since dead cells are excluded from the count.

To simplify the modeling, we assume that bacteria may switch phenotypes only at the time of division [28] and only between two phenotypes, a reasonable assumption for ciprofloxacin. Depending on how the phenotype of the mother cell is inherited by the daughter cells at division, we propose two variants of the model and for each of them, we compute the cell probability distribution which can be compared to experimental data.

The first model is sketched in Fig. 1A. Let ***n*** = (*n*_1_, …, *n*_*P*_) be the population vector of the *P* different cell phenotypes and *p*(***n***, *t, c*) the probability to observe that configuration at time *t* when the concentration of antibiotics is *c*. Each bacterial phenotype *i* is characterized by its growth rate *λ*_*i*_, death rate *ν*_*i*_, and by a transition probability *π*_*i*_ (transition to state *i*). We also assume that a cell division produces two cells, and one of them is identical to the mother cell while the other one is in a different state drawn according to the distribution (*π*_*i*_)1*≤i≤P* .

**FIG. 1:**
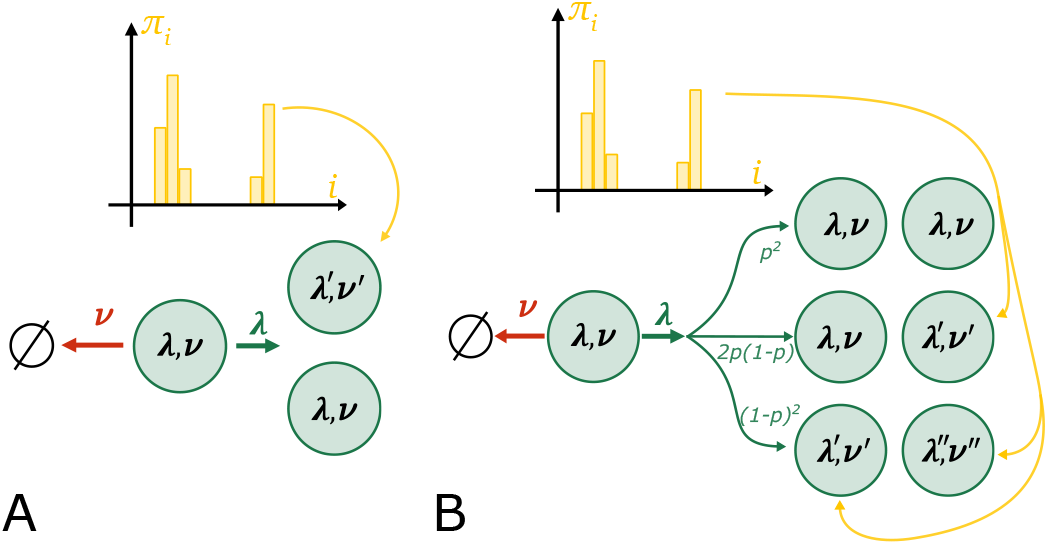
**A** In model 1, a cell may die with rate *ν* or divide with rate *λ*. Then, division produces a cell in the same state and a cell in a new state, drawn from the distribution *π*. **B** In model 2, three outcomes can occur for the two daughter cells following division depending on the heredity probability *p*.

We describe this process at the level of the bacterial population dynamics using a master equation. To write this equation, it is convenient to introduce the vector ***u***_***i***_ = (0, …, 0, 1, 0, …, 0) (of size *P*) where 1 is in position *i*.A transition from state ***n* + *u***_***i***_ at time *t* to state ***n*** at time *t* + *dt* occurs when a cell of the subgroup *i* dies during *dt*, which happens with probability (*n*_*i*_ + 1)*ν*_*i*_*dt*. Alternatively, a transition from state ***n*− *u***_***i***_ to state ***n*** occurs if a cell of the subgroup *j* ≠ *i* divides during *dt* and produces one cell of the subgroup *j* and one cell of the subgroup *i* with probability *π*_*i*_*n*_*j*_*λ*_*j*_*dt*. The assumption of the transmission of the state of the mother cell to one of the daughter cells [29–34] implies mother-daughter correlations [35], which are indeed observed.

From the master equation of the model (see details in Suppl. Mat.), we derive the following equation for the generating function 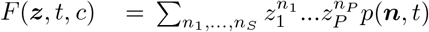:

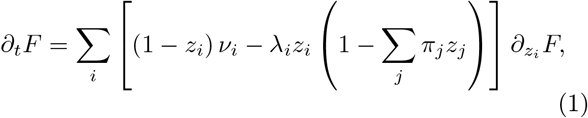

where the first term on the right-side of the equation describes death events, while the second term results from division events.

In practice, the stochastic partition of mRNAs and mother proteins at division influences the phenotypes of daughters cells [36, 37] and leads to variable mother-daughter correlations [34]. To take this variability into account, we modified the previous model and built model 2, also shown in Fig. 1B. The main difference is that now, following division, each daughter cell remains in the mother’s state with probability *p* or acquires a different state drawn from the distribution (*π*_*i*_)_1*≤i≤P*_ with probability 1 −*p*. This has the advantage to make the strength of heredity or equivalently the mother-daughter correlations, tunable and only controlled by *p*. In a further generalization of this second model, one can replace the parameter *p* by an heredity matrix that takes into account variable levels of heredity among all cell types. All these models are detailed in the Suppl. Mat.

We have solved analytically these models using the method of characteristics [38], when there is a separation of timescales in the dynamics. More precisely, we assume that there is a single fast (or active) state while all other states are slow (or dormant), in order to have an analytically solvable system. Without loss of generality, we define the fast active state to be state 1, so that ∀*j* > 1, λ_*j*_ ≪ λ_1_ and ∀*j* > 1, *ν*_*j*_ ≪ *ν*_1_.

For model 1, the analytical solution of Eq. 1 depends on the susceptibility *q*_*i*_ = *ν*_*i*_*/*λ_*i*_, which represents the probability of a cell type *i* to die if *ν*_*i*_ < λ_*i*_ and if only state *i* is considered. We deduce from the generating function, the time evolution of the probabilities *p*_*n*_(*t*|*c*) of observing *n* cells at time *t* given the concentration of antibiotics *c* and the initial condition *n*_0_(*k*) = *n*_*k*_(*t* = 0).

When there is only a single bacterial phenotype (*P* =1), defined by λ_1_ and *ν*_1_, the probability of extinction is given by 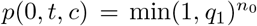 at large times with *n*_0_ = *n*_0_(1). In addition, for *t* ≫1*/*λ_1_,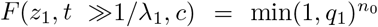 , which is independent of *z*_1_. Thus, the probability to have any finite population in this limit is 0, and at large time, the only possible states with non zero probabilities are the empty state or arbitrarily large populations , as expected from the theory of birth-death processes [39]. To summarize, when there is a single bacterial phenotype and thus a single timescale, no tail can exist in the distribution for small populations. This is illustrated on Fig. 2A, where the probability to acquire state 2 is *π*_2_ = 0, and the system behaves as if there was only one state. As expected, only the extinction probability converges towards a finite value, while the probability to observe a finite number of individuals goes to 0 over the timescale of state 1.

**FIG. 2:**
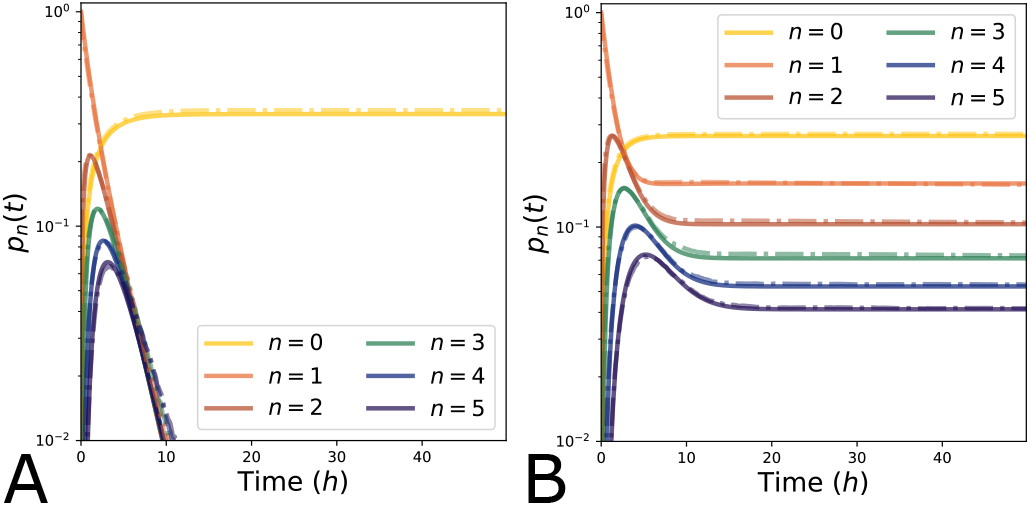
**A** *With a single phenotype:* Time evolution of the probability distribution for different populations starting from one individual in state 1 with *q*_1_ = 1/3, *q*_2_ = 0.1, *π*_1_ = 1.0. Solid lines are analytical expressions, while dotted lines represent Gillespie simulations. **B** *With two phenotypes:* All the other conditions are the same, except for *π*_1_ = 0.3.

The situation changes completely when at least two separate bacterial phenotypes 1, 2 are present, with two different growth rates λ_1_ ≫λ_2_ and death rates *ν*_1_ ≫*ν*_2_. Assuming the initial condition is such that *n*_0_(1) cells are in the first state and none in the second state (*n*_0_(2) = 0), we find the following extinction probability (see Suppl.Mat. for details):

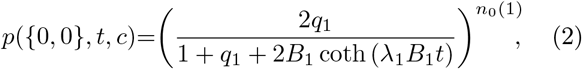

with 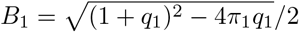.

For the active state, we used the value λ_1_ ∼0.6 *h*^−1^ (bulk growth rate leading to a doubling time of ∼ 1.2 *h*, in practice the single cell growth rate may be larger, at least 1 *h*^−1^ which would simply accelerate the dynamics), with the death rate *ν*_1_ ∼0.2 *h*^−1^. For the *dormant/slowly growing* state, we use λ_2_ ∼ 10^−3^ *h*^−1^ and *ν*_2_ ∼ 10^−4^ *h*^−1^. Our analytical expression is successfully compared to stochastic simulations in Fig. 2B. The important point is that now we observe a plateau at large times (over a timescale 1*/*λ_1_, 1*/v*_1_ ≪*t* ≪1*/*λ_2_, 1*/v*_2_), confirming the experimental observation that the probability to obtain small finite populations can be significant.

Let us now consider model 2. For symmetric division, we can compute the evolution of the distribution for different values of *p*. In particular the extinction probability in this case depends on 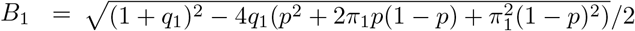 and we obtain the same equation Eq. 2 with this modified *B*_1_ for the extinction probability.

The dependency on heredity (through the parameter *p*) is illustrated in Figs. 3 A and B for the probabilities *p*_*n*_(*t*). We observe that *p* strongly modifies the evolution of the distributions, but does not modify the survival probability significantly. It mostly affects the time to reach a steady state, since increasing *p* increases the timescale over which the distribution stabilizes. In other words, the initial state acquires a larger “inertia” when heredity gets stronger. Here, heredity increases early amplification of the distributions, leading *p*_*n*_(*t*) to reach a maximum before decreasing. Indeed, heredity increases the fraction of cells in state 1 here (because we assume that the initial cell is in state 1), thus favoring a growing population (because more cells are active).

**FIG. 3:**
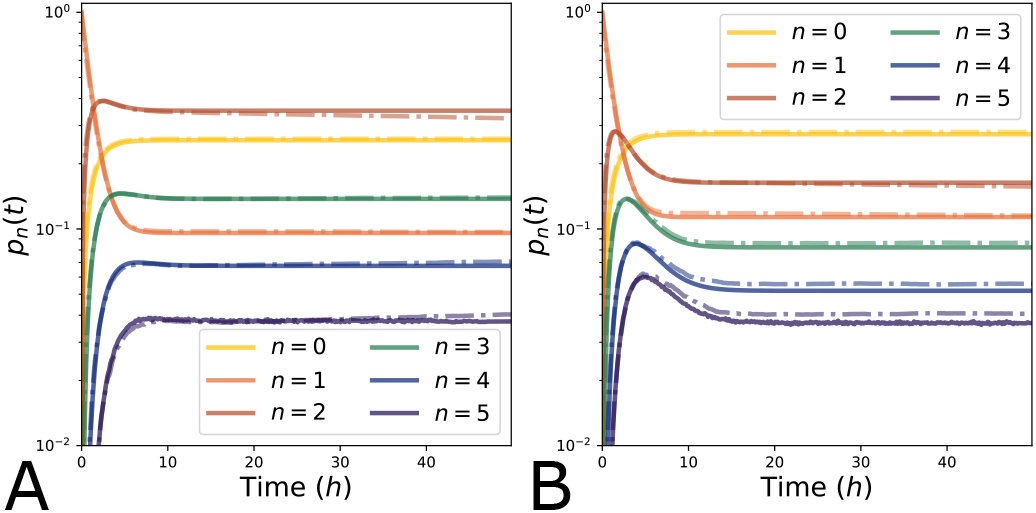
**A** Time evolution of the cell distribution starting from one cell in state 1 with *q*_1_ = 1*/*3, *q*_2_ = 0.1, *π*_1_ = 0.3 and *p* = 0.1, corresponding to a weak heredity. **B** Same plot and conditions except for *p* = 0.5, which represents a significant heredity.Solid lines are analytical expressions, while dotted lines represent Gillespie simulations.

We now compare our model to experiments. In practice, the inoculation size of droplets is stochastic, which we can model by assuming that the content of droplets (in terms of number and cell types) is initially drawn according to a Poisson binomial law with parameters *µ* and *π*_1_. This process accounts first for the number *n*_0_ of individuals in the droplet, and then for its composition in terms of cell types, resulting in the product of a Poisson distribution with parameter *µ* and a binomial distribution of parameters *π*_1_ and *n*_0_ conditioned on *n*_0_ (see Suppl. Mat.). The parameter *µ* is well known from experiments using the cell distribution at *t* = 0. We find that the extinction probability is given by

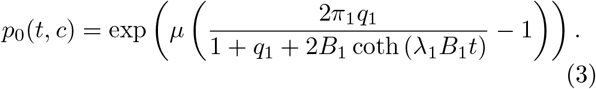

We have obtained a complementary expression for the probability of having one cell (see Suppl. Mat.) and we have used the values taken by these two probabilities at large times to infer the parameters *π*_1_, *q*_1_ and *p* (see Suppl. Mat. for tables summarizing the parameters) from experimental data for ciprofloxacin taken from Ref. [24]. In practice, large times means a duration of 24 *h*, which is indeed large as compared to the inverse growth rate (λ_1_∼ 0.6 *h*^−1^ for the bulk growth rate) but small as compared to the timescale of dormant/inhibited states. In Fig. 4A, we compare the population distributions predicted by our analytical theory with experimental measurements. In the future, we hope to be able to further test of this model using data at finite time.

**FIG. 4:**
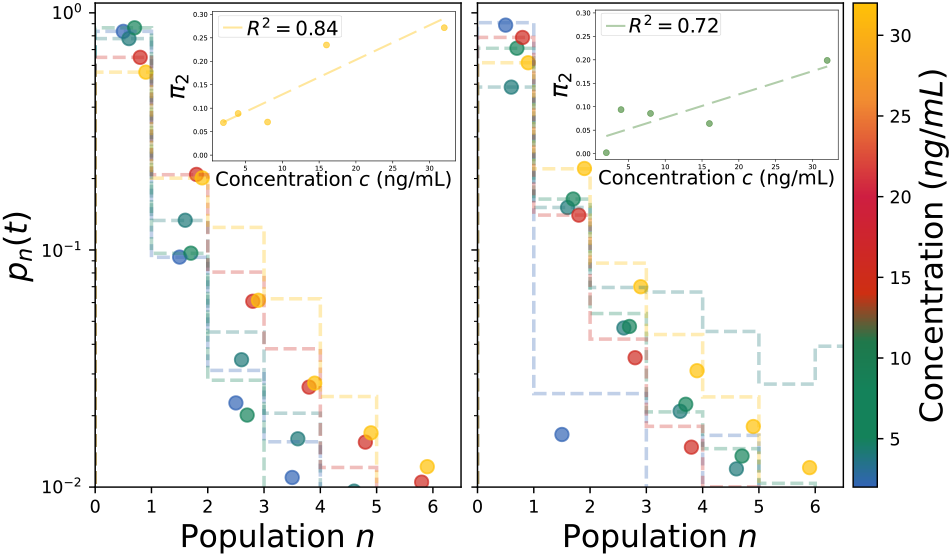
Comparison of the model to experiments with ciprofloxacin. Cell distributions are shown after 24 *h* for two repetitions of the same experimental setting. The dotted lines are the experimental data of cell numbers measured inside droplets while bullet points are analytic predictions of the model using inferred parameters. The insets display linear regressions of *π*_2_ = 1− *π*_1_ (transition probability from state 1 towards state 2).We find that *π*_2_ increases with antibiotic concentration.

An important question concerns the transition between dormant/slow growth and the active state. If the transition towards dormant/slow growth state is induced by the antibiotics, we expect that this rate should increase with the antibiotic concentration. Growth rates are also dependent on antibiotic concentration [3]. To address this question, we analyzed experimental data corresponding to an antibiotic concentration, which is small with respect to the half inhibitory concentration (*c*≪ *IC*_50_). In this regime, the simplest choice for the dependency of *π*_1_, *π*_2_ on drug concentration *c* is a linear dependence (first order expansion). This dependence is well supported by the data as shown in Fig. 4B (see Suppl. Mat. for details). Moreover, *q*_1_ is not well described by a linear dependence, and there is no clear trend appearing for *q*_1_ as a function of the drug concentration. This is expected because both the death and division rates depend on the antibiotic concentration.

Experimentally, to measure the single-cell antibiotic susceptibility (at the single individual level) [23, 24, 40], a threshold in the final number of observed bacteria is typically fixed below which it is considered that the droplet is “dead”. However, the frequency of “dead” droplets is not equal to the extinction probability, in particular for bistable bacterial populations where a small number of bacteria (below the threshold) may be explained by the presence of dormant individuals. Our model gives an interpretation for droplets where the final number of bacteria is below the threshold, and allows to estimate the parameters (growth rate, death rate and transition rate) from the distribution of populations.

To summarize, this work shows that growth heterogeneity in small bacterial population exposed to antibiotics can be modeled using discrete phenotype models. Further, the observed linear dependence of the transition probability into the dormant/slow growth state supports the view that such a state is probably induced by the antibiotics in this experiment rather than resulting from slowly growing cells before antibiotic exposure [22]. Interestingly, a recent study based on fluctuation tests found that the transition to persistence in certain cancers is, as in our case, mainly drug-induced and dose-dependent [41]. While our model has been built to explain the response of bacteria to ciprofloxacin, it is agnostic about the cause of growth bistability, thus, it could be extended to other antibiotics and even to situations in which the growth bistability is triggered by other forms of stress, such as that caused by a change of medium quality [42] or by the application of a combination of antibiotics [19, 43].

## ACKNOWLEDGMENTS

We acknowledge key discussions with C. Baroud and E. Maikranz, which motivated this work, and further insightful discussions with M. El Karoui.

## References

[1] P. Greulich, M. Scott, M. R. Evans, and R. J. Allen. Growth-dependent bacterial susceptibility to ribosome-targeting antibiotics. Mol. Syst. Biol., 11(3):796, 2015.

[2] P. Greulich, J. Doležal, M. Scott, M. R. Evans, and R. J. Allen. Predicting the dynamics of bacterial growth inhibition by ribosome-targeting antibiotics. Phys. Biol., 14:065005, 2017.

[3] B. Ledoux and D. Lacoste. Inhibition of bacterial growth by antibiotics: a minimal model. Phys. Biol., 22(6):066007, 2025.

[4] M. Kals, E. Kals, J. Kotar, D. Allen, L. Mancini, and C. Pietro. Antibiotics change the population growth rate heterogeneity and morphology of bacteria. PLoS Pathog., 21(2):e1012924, 2025.

[5] C. J. Skalnik, S. Y. Cheah, M. Y. Yang, M. B. Wolff, R. K. Spangler, L. Talman, J. H. Morrison, S. M. Peirce, E. Agmon, and M. W. Covert. Whole-cell modeling of E. coli colonies enables quantification of single-cell heterogeneity in antibiotic responses. PLoS Comput. Biol., 19(6):e1011232, 2023.

[6] H. Zhu, Y. Xiong, Z. Jiang, Q. Liu, and J. Wang. Quantifying Dynamic Phenotypic Heterogeneity in Resistant Escherichia coli under Translation-Inhibiting Antibiotics. Adv. Sci. (Weinh.), 11(11):2304548, 2024.

[7] J. B. Deris, M. Kim, Z. Zhang, H. Okano, R. Hermsen, A. Groisman, and T. Hwa. The Innate Growth Bistability and Fitness Landscapes of Antibiotic-Resistant Bacteria. Science, 342(6162):1237435, 2013.

[8] A. Roy and S. Klumpp. Simulating Genetic Circuits in Bacterial Populations with Growth Heterogeneity. Biophys. J., 114(2):484–492, 2018.

[9] R. Malka, E. Shochat, and V. Rom-Kedar. Bistability and Bacterial Infections. PLoS One, 5:e10010, 2010.

[10] N. Frenkel, R. Saar Dover, E. Titon, Y. Shai, and V. Rom-Kedar. Bistable Bacterial Growth Dynamics in the Presence of Antimicrobial Agents. Antibiotics, 10:87, 2021.

[11] J. Karslake, J. Maltas, P. Brumm, and K. B. Wood. Population Density Modulates Drug Inhibition and Gives Rise to Potential Bistability of Treatment Outcomes for Bacterial Infections. PLoS Comput. Biol., 12:e1005098, 2016.

[12] S. Ghosh, K. Sureka, B. Ghosh, I. Bose, J. Basu, and M. Kundu. Phenotypic heterogeneity in mycobacterial stringent response. BMC Syst. Biol., 5:18, 2011.

[13] H. J. Hindley, Z. Gong, S. Moradian, M. G. Giuliano, A. Sapelkin, I. Kotta-Loizou, M. Buck, C. Engl, and A. Y. Weiße. Heterogeneity in responses to ribosome-targeting antibiotics mediated by bacterial RNA repair. Nat. Commun., 16:9620, 2025.

[14] M. S. Svenningsen and N. Mitarai. Simple bacterial growth model for the formation of spontaneous and triggered dormant subpopulations. Phys. Rev. Res., 6(3):033072, 2024.

[15] S. Jaramillo-Riveri, J. Broughton, A. McVey, T. Pilizota, M. Scott, and M. El Karoui. Growth-dependent heterogeneity in the DNA damage response in Escherichia coli. Mol. Syst. Biol., 18(5):MSB202110441, 2022.

[16] P. Patra and S. Klumpp. Population Dynamics of Bacterial Persistence. PLoS One, 8(5):e62814, 2013.

[17] M. Stevanovic, J. P. Teuber Carvalho, P. Bittihn, and D. Schultz. Dynamical model of antibiotic responses linking expression of resistance genes to metabolism explains emergence of heterogeneity during drug exposures. Phys. Biol., 21:036002, 2024.

[18] M. Arnoldini, I. A. Vizcarra, R. Peña-Miller, N. Stocker, M. Diard, V. Vogel, R. E. Beardmore, W.-D. Hardt, and M. Ackermann. Bistable Expression of Virulence Genes in Salmonella Leads to the Formation of an Antibiotic-Tolerant Subpopulation. PLoS Biol., 12(8):e1001928, 2014.

[19] J. Broughton, A. Fraisse, and M. El Karoui. Suppression of bacterial cell death underlies the antagonistic interaction between ciprofloxacin and tetracycline. Mol. Syst. Biol., 22(1):102–118, 2026.

[20] T. Dörr, M. Vulić, and K. Lewis. Ciprofloxacin causes persister formation by inducing the TisB toxin in Escherichia coli. PLoS Biol., 8(2):e1000317, 2010.

[21] T. Dörr, K. Lewis, and M. Vulić. SOS response induces persistence to fluoroquinolones in Escherichia coli. PLoS Genet., 5(12):e1000760, 2009.

[22] M. Umetani, M. Fujisawa, R. Okura, T. Nozoe, S. Suenaga, H. Nakaoka, E. Kussell, and Y. Wakamoto. Observation of persister cell histories reveals diverse modes of survival in antibiotic persistence. eLife, 14:e79517, 2025.

[23] L. Le Quellec, A. Aristov, S. Gutiérrez Ramos, G. Amselem, J. Bos, Z. Baharoglu, D. Mazel, and C. N. Baroud. Measuring single-cell susceptibility to antibiotics within monoclonal bacterial populations. PLoS One, 19(8):e0303630, 2024.

[24] E. Maikranz, A. Aristov, L. Le Quellec, and C. Baroud. Extending digital biology: bacterial survival and morphological heterogeneity under antibiotic stress, 2025.

[25] G. Amselem, C. Guermonprez, B. Drogue, S. Michelin, and C. N. Baroud. Universal microfluidic platform for bioassays in anchored droplets. Lab Chip, 16(21):4200–4211, 2016.

[26] A. Barizien, M. S. Suryateja Jammalamadaka, G. Amselem, and C. N. Baroud. Growing from a few cells: combined effects of initial stochasticity and cell-to-cell variability. J. R. Soc. Interface, 16(153):20180935, 2019.

[27] K. M. T. Rahman and N. C. Butzin. Counter-on-chip for bacterial cell quantification, growth, and live-dead estimations. Sci. Rep., 14(1):782, 2024.

[28] L. Brettner and K. Geiler-Samerotte. Single-cell heterogeneity and the growth laws. bioRxiv, page 2024.04.19.590370, 2024.

[29] D. Huh and J. Paulsson. Non-genetic heterogeneity from stochastic partitioning at cell division. Nat. Genet., 43:95–100, 2011.

[30] D. Huh and J. Paulsson. Random partitioning of molecules at cell division. Proc. Natl. Acad. Sci. U.S.A., 108:15004–15009, 2011.

[31] Q. Chai, B. Singh, K. Peisker, N. Metzendorf, X. Ge, S. Dasgupta, and S. Sanyal. Organization of Ribosomes and Nucleoids in Escherichia coli Cells during Growth and in Quiescence *. J. Biol. Chem., 289:11342–11352, 2014.

[32] A. Papagiannakis, Q. Yu, S. K. Govers, W. Lin, N. S. Wingreen, and C. Jacobs-Wagner. Nonequilibrium polysome dynamics promote chromosome segregation and its coupling to cell growth in Escherichia coli. eLife, 14:RP104276, 2025.

[33] A. M. Miangolarra, S. H. Li, J. Joanny, N. S. Wingreen, and M. Castellana. Steric interactions and out-of-equilibrium processes control the internal organization of bacteria. Proc. Natl. Acad. Sci. U.S.A., 118(43):e2106014118, 2021.

[34] J. Lin and A. Amir. From single-cell variability to population growth. Phys. Rev. E, 101:012401, 2020.

[35] J. Grilli, C. Cadart, G. Micali, M. Osella, and M. Cosentino Lagomarsino. The Empirical Fluctuation Pattern of E. coli Division Control. Front. Microbiol., Volume 9-2018, 2018.

[36] D. Antunes and A. Singh. Quantifying gene expression variability arising from randomness in cell division times. J. Math. Biol., 71(2):437–463, 2015.

[37] B. Cerulus, A. M. New, K. Pougach, and K. J. Verstrepen. Noise and Epigenetic Inheritance of Single-Cell Division Times Influence Population Fitness. Curr. Biol., 26(9):1138–1147, 2016.

[38] P. J. Olver. Introduction to partial differential equations, volume 1. Springer, 2014.

[39] D. G. Kendall. On the Generalized “Birth-and-Death” Process. Ann. Math. Stat., 19(1):1–15, 1948.

[40] A. Samimi, N. Verdon, R. J. Allen, and M. A. Rosenbaum. Probing Antibiotic Inhibition in Small Bacterial Populations With Combinatorial Droplet Microfluidics. Small Sci., 6(1):e202500421, 2026.

[41] M. Russo, S. Pompei, A. Sogari, M. Corigliano, G. Crisafulli, A. Puliafito, S. Lamba, J. Erriquez, A. Bertotti, M. Gherardi, F. Di Nicolantonio, A. Bardelli, and M. Cosentino Lagomarsino. A modified fluctuation-test framework characterizes the population dynamics and mutation rate of colorectal cancer persister cells. Nat. Genet., 54:976–984, 2022.

[42] O. Kotte, B. Volkmer, J. L. Radzikowski, and M. Heinemann. Phenotypic bistability in Escherichia coli’s central carbon metabolism. Mol. Syst. Biol., 10(7):736, 2014.

[43] R. Chait, A. Craney, and R. Kishony. Antibiotic interactions that select against resistance. Nature, 446(7136):668–671, 2007.

