## Supplementary Materials for "Growth bistability in small bacterial populations exposed to antibiotics"

### Supplementary material for "Growth bistability in small bacterial populations exposed to antibiotics"

B. Ledoux, D. Lacoste

May 21, 2026

#### 1 Small populations under antibiotic stress

The experimental setup we want to model is made of a large number ($N_d > 10^2$) of identical droplets containing nutrients and antibiotics at a controlled concentration $c$ [1, 2, 3]. Each droplet is then distributed with a small number of bacteria (*E.Coli*), which we model by an initial distribution of $n$ (typically a Poisson [4, 5] or negative binomial distribution). This initial distribution has a mean of order 1. After the initialization, the droplets evolve independently for a fixed duration, and the number of individuals in each droplet is monitored. The random variable $n^{(k)}(t, c)$ thus represents the number of individuals at time $t$ after the beginning of the experiment ($t = 0$ corresponds to the initialization) in the droplet $k$, in a medium with antibiotic concentration $c$. Using the large number of independent repetitions of the same experiment (every droplet correspond to a repetition, with different initial conditions), we can study the statistics of the population at time $t$ and antibiotic concentration $c$, and estimate the underlying probability distribution.

Below, we compute this probability distribution in the presence of growth heterogeneity in a theoretical model, which is then simulated with Gillespie's algorithm [6].

##### 1.1 Distribution of bacterial states in a birth/death model with discrete populations

The first model we consider is defined in the main text. We define $p(\boldsymbol{n}, t, c) = p((n_1, ..., n_P), t, c)$ where $P$ is the number of different bacterial states (characterized by their growth rates $(\lambda_i)_{1\leq i\leq P}$ and death rates $(\nu_i)_{1\leq i\leq P}$) and $c$ is the concentration of antibiotics (which we assume to be homogeneous). We introduce $\boldsymbol{u_i} = (0, ..., 0, 1, 0, ..., 0)$ where 1 is in position $i$. The different possible steps leading to state $\boldsymbol{n}$ at $t + dt$ are

- State $\boldsymbol{n} + \boldsymbol{u_i}$ at $t$, a cell of the subgroup $i$ dies during $dt$ (with probability $(n_i + 1)\nu_i dt$)
- State $\boldsymbol{n} - \boldsymbol{u_i}$ at $t$, a cell of the subgroup $j$ divides during $dt$ and gives one cell of the subgroup $j$ and one cell of the subgroup $i$ (with probability $\pi_i n_j \lambda_j dt$). This strong hypothesis is based on the fact that cell division is rather heterogeneous [7, 8, 9, 10, 11, 12]. Therefore we translate the asymmetry of ribosome contents at division to this hypothesis that only one of the bacterial state is drawn depending on the distribution of $\lambda$. This assumption also allows to model inheritance during cell division [13], as correlations are typically observed.
- State $\boldsymbol{n}$ at $t$, and nothing happens (no birth nor death event).

Therefore we have

$$\begin{aligned} p(\boldsymbol{n}, t+dt, c) - p(\boldsymbol{n}, t, c) = &\sum_i \nu_i \left[ (n_i+1) p(\boldsymbol{n}+\boldsymbol{u_i}, t, c) - n_i p(\boldsymbol{n}, t, c) \right] dt \\ &+ \sum_{i,j} \pi_i \lambda_j \left[ (n_j - \delta_{i,j}) p(\boldsymbol{n}-\boldsymbol{u_i}, t, c) - n_j p(\boldsymbol{n}, t, c) \right] dt. \end{aligned} \tag{1}$$

This yields

$$\partial_t p(\boldsymbol{n}, t, c) = \sum_i \nu_i \left[ (n_i+1) p(\boldsymbol{n}+\boldsymbol{u_i}, t, c) - n_i p(\boldsymbol{n}, t, c) \right] + \sum_{i,j} \pi_i \lambda_j \left[ (n_j - \delta_{i,j}) p(\boldsymbol{n}-\boldsymbol{u_i}, t, c) - n_j p(\boldsymbol{n}, t, c) \right]. \tag{2}$$

Now we introduce the generating function $F(\boldsymbol{z}, t, c) = \sum_{n_1,\dots,n_P} z_1^{n_1} \dots z_P^{n_P} p(\boldsymbol{n}, t) = \sum_{n_1,\dots,n_P} \prod_{k=1}^P z_k^{n_k} p(\boldsymbol{n}, t, c)$. We have that

$$\begin{aligned} \partial_t \sum_{n_1,\dots,n_P} z_1^{n_1} \dots z_P^{n_P} p(\boldsymbol{n}, t, c) = &\sum_{n_1,\dots,n_P} z_1^{n_1} \dots z_P^{n_P} \sum_i \nu_i \left[ (n_i+1) p(\boldsymbol{n}+\boldsymbol{u_i}, t, c) - n_i p(\boldsymbol{n}, t, c) \right] \\ &+ \sum_{n_1,\dots,n_P} z_1^{n_1} \dots z_P^{n_P} \sum_{i,j} \pi_i \lambda_j \left[ (n_j - \delta_{i,j}) p(\boldsymbol{n}-\boldsymbol{u_i}, t, c) - n_j p(\boldsymbol{n}, t, c) \right] \\ = &\sum_i \nu_i \left[ \sum_{n_1,\dots,n_P} \prod_{k \neq i}^P z_k^{n_k} (n_i+1) z_i^{n_i} p(\boldsymbol{n}+\boldsymbol{u_i}, t, c) - z_i \sum_{n_1,\dots,n_P} \prod_{k \neq i}^P z_k^{n_k} n_i z_i^{n_i - 1} p(\boldsymbol{n}, t, c) \right] \\ &+ \sum_{i,j} \pi_i \lambda_j \left[ \sum_{n_1,\dots,n_P} \prod_{k \neq j}^P z_k^{n_k} (n_j - \delta_{i,j}) z_j^{n_j} p(\boldsymbol{n}-\boldsymbol{u_i}, t, c) - z_j \sum_{n_1,\dots,n_P} \prod_{k \neq j}^P z_k^{n_k} n_j z_j^{n_j - 1} p(\boldsymbol{n}, t, c) \right] \\ = &\sum_i \nu_i (1 - z_i) \partial_{z_i} F(\boldsymbol{z}, t, c) + \sum_{i,j} \pi_i \lambda_j z_j (z_i - 1) \partial_{z_j} F(\boldsymbol{z}, t, c). \end{aligned} \tag{3}$$

So that we find the partial differential equation

$$\partial_t F(\boldsymbol{z}, t, c) = \sum_i (1 - z_i) \left[ \nu_i \partial_{z_i} F(\boldsymbol{z}, t, c) - \pi_i \left( \sum_j \lambda_j z_j \partial_{z_j} F(\boldsymbol{z}, t, c) \right) \right], \tag{4}$$

or equivalently

$$\partial_t F(\boldsymbol{z}, t, c) = \sum_i \left[ (1 - z_i) \nu_i - \lambda_i z_i \left( 1 - \sum_j \pi_j z_j \right) \right] \partial_{z_i} F(\boldsymbol{z}, t, c). \tag{5}$$

We use the method of characteristics and we look for a set of variables $u, v_1, \dots v_P$ such that

$$\frac{dF}{du} = \frac{dt}{du} \partial_t F + \sum_i \frac{dz_i}{du} \partial_{z_i} F, \tag{6}$$

and

$$\begin{aligned}
&\frac{dF}{du} = 0 \\
&\frac{dt}{du} = 1 \\
\forall i,\ &\frac{dz_i}{du} = -\left(\pi_i \lambda_i z_i^2 - z_i\left(\lambda_i + \nu_i - \lambda_i \sum_{j\neq i} \pi_j z_j\right) + \nu_i\right).
\end{aligned} \tag{7}$$

With the initial parametric conditions

$$\begin{aligned}
t(u = 0, \{v_i\}_i) &= 0 \\
z_i(u = 0, \{v_j\}_j) &= v_i \\
F(\{v_i\}_i, u = 0, c) &= \prod_{k=1}^{P} v_k^{n_0(k)},
\end{aligned} \tag{8}$$

which correspond to the initial condition $p(n, t = 0) = \prod_{k=1}^{P} \delta_{n_k, n_0(k)}$. Therefore we obtain coupled quadratic first order differential equations for $z_i$.

**Separation of timescales** Now we assume that all states except one are dormant states. The fast metabolic state is state 1, so that $\forall j > 1, \lambda_j \ll \lambda_1$ and $\forall j > 1, \nu_j \ll \nu_1$. In this case, we can consider that over the typical timescale of $z_1$, all $z_j$ are constant. In this case

$$\begin{aligned}
&\frac{dF}{du} = 0 \\
&\frac{dt}{du} = 1 \\
&du = -\frac{dz_1}{\pi_1 \lambda_1 z_1^2 - (\nu_1 + \lambda_1(1 - \sum_{j>1} \pi_j z_j))z_1 + \nu_1} \\
\forall j >1,\ &\frac{dz_j}{du} = 0.
\end{aligned} \tag{9}$$

Defining $\Delta_1(\{z_j\}_{j\neq i}) = \lambda_1^2 \left[\left(1 + \frac{\nu_1}{\lambda_1} - \sum_{j>1} \pi_j z_j\right)^2 - 4\frac{\pi_1 \nu_1}{\lambda_1}\right]$, we obtain

$$du = -\frac{dz_1}{\pi_1 \lambda_1 \left(z_1 - \frac{(\nu_1 + \lambda_1(1-\sum_{j>1}\pi_j z_j))}{2\pi_1\lambda_1} + \frac{\sqrt{\Delta_1(\{z_j\}_{j>1})}}{2\pi_1\lambda_1}\right)\left(z_1 - \frac{(\nu_1 + \lambda_1(1-\sum_{j>1}\pi_j z_j))}{2\pi_1\lambda_1} - \frac{\sqrt{\Delta_1(\{z_j\}_{j>1})}}{2\pi_1\lambda_1}\right)}. \tag{10}$$

Therefore, if we define $A_1(\{z_j\}_{j>1}) = \frac{(\nu_1 + \lambda_1(1-\sum_{j>1}\pi_j z_j))}{2\lambda_1}$, $B_1(\{z_j\}_{j>1}) = \frac{\sqrt{\Delta_1(\{z_j\}_{j>1})}}{2\lambda_1}$, we find

$$du = -\frac{dz_1}{2\lambda_1 B_1}\left(\frac{1}{\pi_1 z_1 - A_1 - B_1} - \frac{1}{\pi_1 z_1 - A_1 + B_1}\right), \tag{11}$$

and thus by integrating the last equation using the initial condition $v_1 = z_1$ at $t = 0$,

$$
\begin{aligned}
F(\{v_i\}_i, u, c) &= \prod_{k=1}^{P} v_k^{n_0(k)} \\
t &= u \\
\forall j > 1,\ z_j &= v_j \\
-2\pi_1\lambda_1 B_1 u &= \ln\left(\frac{\pi_1 z_1 - A_1 - B_1}{\pi_1 z_1 - A_1 + B_1}\frac{\pi_1 v_1 - A_1 + B_1}{\pi_1 v_1 - A_1 - B_1}\right).
\end{aligned}
\tag{12}
$$

Therefore we get

$$
\begin{aligned}
&\forall j > 1, z_j = v_j \\
&v_1 = \frac{(A_1 - B_1)(A_1 + B_1 - \pi_1 z_1)e^{2\pi_1\lambda_1 B_1 u} - (A_1 + B_1)(A_1 - B_1 - \pi_1 z_1)}{\pi_1(A_1 + B_1 - z_1)e^{2\pi_1\lambda_1 B_1 u} - \pi_1(A_1 - B_1 - z_1)} \\
&\quad = \frac{z_1\left(\frac{A_1}{B_1} - \coth(\lambda_1 B_1 t)\right) - \frac{B_1}{\pi_1}\left(\frac{A_1^2}{B_1^2} - 1\right)}{\frac{\pi_1 z_1}{B_1} - \frac{A_1}{B_1} - \coth(\lambda_1 B_1 t)}.
\end{aligned}
\tag{13}
$$

We introduce $q_i = \frac{\nu_i}{\lambda_i}$ which is the extinction probability for $\nu_i < \lambda_i$ (else this probability is 1), then

$$
\begin{aligned}
A_1(\{z_j\}_{j\neq i}) &= \frac{(1 + q_1 - \sum_{j>1}\pi_j z_j)}{2} \\
B_1(\{z_j\}_{j\neq i}) &= \frac{1}{2}\sqrt{\left(1 + q_1 - \sum_{j>1}\pi_j z_j\right)^2 - 4\pi_1 q_1},
\end{aligned}
\tag{14}
$$

and the generating function is

$$
F(\{z_i\}, t, c) = \left(\frac{z_1\left(\coth(\lambda_1 B_1 t) - \sqrt{1 + \frac{\pi_1 q_1}{B_1^2}}\right) + \frac{q_1}{B_1}}{\coth(\lambda_1 B_1 t) + \sqrt{1 + \frac{\pi_1 q_1}{B_1^2}} - \frac{\pi_1 z_1}{B_1}}\right)^{n_0(1)} \prod_{j>1} z_j^{n_0(j)}. \tag{15}
$$

It is easy to check the normalization, as $\forall i, \forall t, v_i(\{1, ..., 1\}, t) = 1$, we obtain

$$
\forall t, F(\{1, ..., 1\}, t, c) = 1. \tag{16}
$$

From the expression of $F$, we can find the values of the probability of having $n$ individuals at time $t$

$$
p_n(t, c) = \frac{1}{n!}\sum_{k_1+...+k_P=n}\frac{n!}{k_1!...k_P!}\partial_{z_1}^{k_1}...\partial_{z_P}^{k_P} F|_{z_1,...,z_P=0,...,0}, \tag{17}
$$

in particular with two states 1 and 2, if we define

$$F_1 = \frac{z_1\left(\coth\left(\lambda_1 B_1 t\right) - \sqrt{1+\frac{\pi_1 q_1}{B_1^2}}\right) + \frac{q_1}{B_1}}{\coth\left(\lambda_1 B_1 t\right) + \sqrt{1+\frac{\pi_1 q_1}{B_1^2}} - \frac{\pi_1 z_1}{B_1}}, \tag{18}$$

we find that

$$p_n(t,c) = \sum_{k=0}^{n} \frac{\delta_{n_0(2),n-k}}{k!} \frac{n_0(1)!}{(n_0(1)-k)!} F_1^{n_0(1)-k} \sum_{k_1+\ldots+k_P=k} \frac{k!}{k_1!\ldots k_P!} \partial_{z_1}^{k_1}\ldots\partial_{z_P}^{k_P} F_1|_{z_1,\ldots,z_P=0,\ldots,0}. \tag{19}$$

##### 1.1.1 Single state

If there is only a single bacterial state 1 (with growth rate $\lambda_1$ and death rate $\nu_1$), $\pi_1 = 1$, with $r_1 = \lambda_1 - \nu_1$ we get

$$v_1 = \frac{z_1\left(\coth\left(\lambda_1|1-q_1|t\right) - \sqrt{1+\frac{4q_1}{(1-q_1)^2}}\right) + \frac{2q_1}{|1-q_1|}}{\coth\left(\lambda_1|1-q_1|t\right) + \sqrt{1+\frac{4q_1}{(1-q_1)^2}} - \frac{2z_1}{|1-q_1|}}, \tag{20}$$

and thus

$$F(z_1,t,c) = \left(\frac{z_1\left(\coth\left(\lambda_1|1-q_1|t\right) - \sqrt{1+\frac{4q_1}{(1-q_1)^2}}\right) + \frac{2q_1}{|1-q_1|}}{\coth\left(\lambda_1|1-q_1|t\right) + \sqrt{1+\frac{4q_1}{(1-q_1)^2}} - \frac{2z_1}{|1-q_1|}}\right)^{n_0}. \tag{21}$$

Therefore, the probability of extinction in this case is

$$p(0,t,c) = \left(\frac{2q_1}{|1-q_1|\coth\left(\lambda_1|1-q_1|t\right) + |1+q_1|}\right)^{n_0}, \tag{22}$$

which is $p_{ext} = \min(1,q_1)^{n_0}$ at large times. In addition, for $t \to \infty$

$$F(z_1, t\to\infty, c) = \left(\frac{q_1 - z_1\min(1,q_1)}{1+q_1-\min(1,q_1)-z_1}\right)^{n_0} = p_{ext}, \tag{23}$$

so that the probability to have any finite population is 0, and becomes negligible on the division timescale. In practice this means that at large time, the only possible states (with non zero probabilities) are extinct or arbitrarily large populations. This explains why we do not expect any tail close to 0 in the distribution for a single bacterial state.

##### 1.1.2 Two states with separation of timescales

With two separate bacterial states 1, 2 associated to two different growth rates $\lambda_1 \gg \lambda_2$ and death rates $\nu_1 \gg \nu_2$ we have

$$\begin{aligned} v_1 &= \frac{z_1\left(\coth\left(\lambda_1 B_1 t\right) - \sqrt{1+\frac{\pi_1 q_1}{B_1^2}}\right) + \frac{q_1}{B_1}}{\coth\left(\lambda_1 B_1 t\right) + \sqrt{1+\frac{\pi_1 q_1}{B_1^2}} - \frac{\pi_1 z_1}{B_1}} \\ v_2 &= z_2. \end{aligned} \tag{24}$$

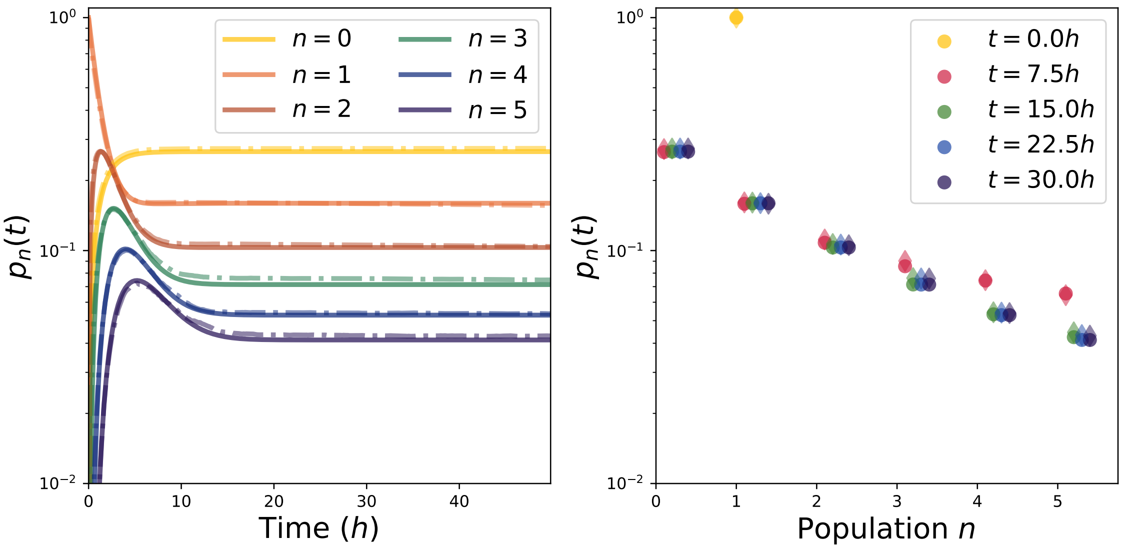


Figure 1: **A** Time evolution of the probabilities for different populations starting from two individuals with $q_1 = 1/3$, $q_2 = 0.1$, $\pi_1 = 0.3$. In dotted lines we show the results of Gillespie simulations. **B** Theoretical histogram for the probability of the number of cells at different times starting from two individuals with $q_1 = 1/3$, $q_2 = 0.1$, $\pi_1 = 0.3$. In diamonds we show the results of Gillespie simulations.

From this we obtain $F$ as

$$F(z_1, z_2, t, c) = \left( \frac{z_1 \left( \coth(\lambda_1 B_1 t) - \sqrt{1 + \frac{\pi_1 q_1}{B_1^2}} \right) + \frac{q_1}{B_1}}{\coth(\lambda_1 B_1 t) + \sqrt{1 + \frac{\pi_1 q_1}{B_1^2}} - \frac{\pi_1 z_1}{B_1}} \right)^{n_0(1)} z_2^{n_0(2)}, \tag{25}$$

where $n_0(i)$ is the initial number of bacteria in state $i$ and the extinction probability is

$$p(\{0,0\}, t, c) = \left( \frac{2q_1}{1 + q_1 + 2B_1 \coth(\lambda_1 B_1 t)} \right)^{n_0(1)} \delta_{n_0(2),0}. \tag{26}$$

Experimentally [1], $\lambda_1 \sim 0.6h^{-1}$, and we assume that the death rate is of the same order of magnitude $\nu_1 \sim 0.2h^{-1}$. For the *dormant/inhibited* state, we use $\lambda_2 \sim 10^{-3}h^{-1}$ and $\nu_2 \sim 10^{-4}h^{-1}$. We find the results of Fig. 1. In particular, we observe that the probabilities plateau at large times. We also observe tails for small populations. We compare our results to Gillespie simulations (stochastic simulation algorithm).

To justify the large times limit, we have that

$$F(z_1,z_2,t\to\infty,c)=z_2^{n_0(2)}\left(\frac{z_1\left(1-\sqrt{1+\frac{\pi_1 q_1}{B_1^2}}\right)+\frac{q_1}{B_1}}{1+\sqrt{1+\frac{\pi_1 q_1}{B_1^2}}-\frac{\pi_1 z_1}{B_1}}\right)^{n_0(1)}, \tag{27}$$

is indeed independent of time $t$. Therefore, this is enough to assume two different bacterial states to recover this distribution. In particular the extinction probability for large times is

$$p(\{0,0\},t\to\infty,c)=\delta_{n_0(2),0}\left(\frac{B_1}{\pi_1}\left(\sqrt{1+\frac{\pi_1 q_1}{B_1^2}}-1\right)\right)^{n_0(1)}, \tag{28}$$

#### 1.2 Distribution with a large number of droplets

##### 1.2.1 General initial distribution

We introduce $n_0^{(k)}(i)$ the initial number of individuals in state $i$ in cell $k$. If we assume that the content of droplets is initially drawn according to a law $p_{n_0(1),n_0(2)}$ for bimodal bacterial states, under separation of timescales. According to the previous part, we have

$$p_n(t,c|n_0(1),n_0(2))=\sum_{k_1+k_2=n}\frac{1}{k_1!k_2!}\partial_{z_1}^{k_1}\partial_{z_2}^{k_2}\ F(z_1,z_2,t,c|n_0(1),n_0(2))|_{\boldsymbol{z}=\boldsymbol{0}}\,, \tag{29}$$

where $n_0$ is the initial number of individuals in the cell. And thus

$$p_n(t,c)=\sum_{n_0(0),n_1(0)=0,0}^{\infty}p_{n_0(1),n_0(2)}\sum_{k_1+k_2=n}\frac{1}{k_1!k_2!}\partial_{z_1}^{k_1}\partial_{z_2}^{k_2}\ F(z_1,z_2,t,c|n_0(1),n_0(2))|_{z_1,z_2=0,0}\,, \tag{30}$$

From this expression we can compute

$$p_0(t,c)=\sum_{n_0(1),n_0(2)=0,0}^{\infty}p_{n_0(1),n_0(2)}\left(\frac{\pi_1 q_1/B_1}{\sqrt{1+\frac{\pi_1 q_1}{B_1^2}}+\coth(\lambda_1 B_1 t)}\right)^{n_0(1)}\delta_{0,n_0(2)}, \tag{31}$$

and thus

$$\begin{aligned} p_0(0,c) &= p_{0,0}\\ p_0(t\to\infty,c) &= \sum_{n_0(1)=0}^{\infty}\frac{p_{n_0(1),0}}{(2(1-\pi_2))^{n_0(1)}}\left(1+q_1-\sqrt{(1+q_1)^2-4\pi_1 q_1}\right)^{n_0(1)} \end{aligned} \tag{32}$$

Therefore, $p_0(t\to\infty,c)/p_0(0,c)$ is in principle an increasing function of both $q_1$ and $q_2$.

##### 1.2.2 Poisson binomial distribution

We assume in this part that the initial distribution is a Poisson binomial distribution with parameter $\mu$ and $\pi_1$, such that

$$p_{n_0(1),n_0(2)} = \frac{e^{-\mu}\mu^{n_0(1)+n_0(2)}}{n_0(1)!n_0(2)!}\pi_1^{n_0(1)}(1-\pi_1)^{n_0(2)}. \tag{33}$$

Therefore, we can compute explicitly

$$\begin{aligned}
p_0(0,c) &= e^{-\mu} \\
p_0(t,c) &= \sum_{n_0(1)=0}^{\infty} \frac{e^{-\mu}\left(\pi_1\mu\right)^{n_0(1)}}{n_0(1)!}\left(\frac{\pi_1 q_1/B_1}{\sqrt{1+\frac{\pi_1 q_1}{B_1^2}}+\coth\left(\lambda_1 B_1 t\right)}\right)^{n_0(1)} \\
&= \exp\left(\mu\left(\frac{\pi_1 q_1/B_1}{\sqrt{1+\frac{\pi_1 q_1}{B_1^2}}+\coth\left(\lambda_1 B_1 t\right)}-1\right)\right) \\
p_0(t\to\infty,c) &= \exp\left(\mu\left(\frac{\pi_1 q_1/B_1}{\sqrt{1+\frac{\pi_1 q_1}{B_1^2}}+1}-1\right)\right)
\end{aligned} \tag{34}$$

We obtain a distribution with a decreasing tail close to $n=0$ at large time, as shown in Fig. 2. This is indeed similar to what is observed experimentally, with a initially peaked distribution ultimately widening close to 0. Furthermore, the quantity

$$r_{00} = \frac{\ln(p_0(t\to\infty,c))}{\ln(p_0(0,c))} = 1-\frac{\pi_1 q_1/B_1}{\sqrt{1+\frac{\pi_1 q_1}{B_1^2}}+1}, \tag{35}$$

depends on the two parameters $\pi_1, q_1$ and this is equivalent to

$$q_1 = \frac{r_{00}(1-r_{00})}{\pi_1+r_{00}-1}, \tag{36}$$

And

$$\begin{aligned}
\frac{p_1(t,c)}{\mu p_0(t,c)} &= (1-\pi_1)+\pi_1\left(\frac{\coth\left(\lambda_1 B_1 t\right)^2-1}{\left(\coth\left(\lambda_1 B_1 t\right)+\sqrt{1+\frac{\pi_1 q_1}{B_1^2}}\right)^2}\right. \\
&\left.+\frac{\pi_2 q_1(1+q_1)}{4B_1^3}\frac{\coth\left(\lambda_1 B_1 t\right)+\sqrt{1+\frac{\pi_1 q_1}{B_1^2}}+B_1\lambda_1 t(1-\coth\left(\lambda_1 B_1 t\right)^2)-\frac{\pi_1 q_1}{B_1^2}\frac{1}{\sqrt{1+\frac{\pi_1 q_1}{B_1^2}}}}{\left(\coth\left(\lambda_1 B_1 t\right)+\sqrt{1+\frac{\pi_1 q_1}{B_1^2}}\right)^2}\right),
\end{aligned} \tag{37}$$

so that for large times

$$r_{01} = \frac{p_1(t\to\infty,c)}{\mu p_0(t\to\infty,c)} = (1-\pi_1)\left(1+\frac{\pi_1 q_1(1+q_1)}{4B_1^3}\frac{1}{1+\frac{\pi_1 q_1}{B_1^2}+\sqrt{1+\frac{\pi_1 q_1}{B_1^2}}}\right), \tag{38}$$

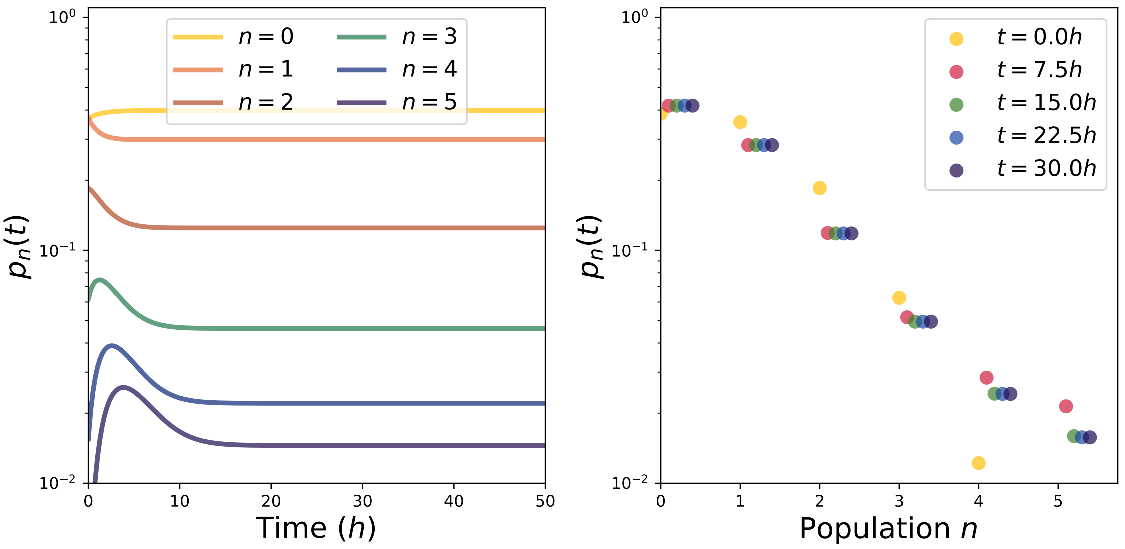


Figure 2: **A** Time evolution of the probabilities for different populations starting from a Poisson binomial distribution with parameters $\mu = 1, q_1 = 1/3$, $q_2 = 0.1$, $\pi_1 = 0.3$. **B** Theoretical histogram for the probability of the number of cells at different times starting from a Poisson distribution of parameter $\mu = 1, q_1 = 1/3$, $q_2 = 0.1$, $\pi_1 = 0.3$.

which also depends only on $\pi_1, q_1$, and is equivalent to

$$q_1 = \frac{B_1}{\pi_1}\left(\frac{r_{01}}{1-\pi_1} - 1\right)(1 + q_1 + 2B_1)\,, \tag{39}$$

from which using the equation on $r_{00}$, we can get a closed equation on $\pi_1$. From these equations we estimate the parameters $\pi_1, q_1$ and $p$ from data for ciprofloxacin [3] and compute the corresponding population distributions. The appearance of subpopulations with different phenotypes under ciprofloxacin stress has been shown [14, 15, 16, 17, 18].

##### 1.2.3 Experimental parameters

The parameters estimated using the methodology presented earlier (based on the estimations of $r_{00}$ and $r_{01}$) from ciprofloxacin experiments are summarized in Table 1.

| $c$ $(ng/mL)$ | $\mu$ | $\pi_1$ | $q_1$ | $c$ $(ng/mL)$ | $\mu$ | $\pi_1$ | $q_1$ |
|---|---|---|---|---|---|---|---|
| 2.0 | 0.67 | 0.93 | 0.99 | 2.0 | 1.29 | 1.00 | 0.95 |
| 4.0 | 0.89 | 0.91 | 1.05 | 4.0 | 1.68 | 0.91 | 0.73 |
| 8.0 | 0.69 | 0.93 | 1.19 | 8.0 | 1.22 | 0.91 | 1.04 |
| 16.0 | 0.89 | 0.76 | 0.99 | 16.0 | 1.15 | 0.94 | 1.15 |
| 32.0 | 0.93 | 0.73 | 0.68 | 32.0 | 1.11 | 0.80 | 1.03 |

Table 1: Estimated parameters for the first (left table) and second (right table) repetitions of the experiments.

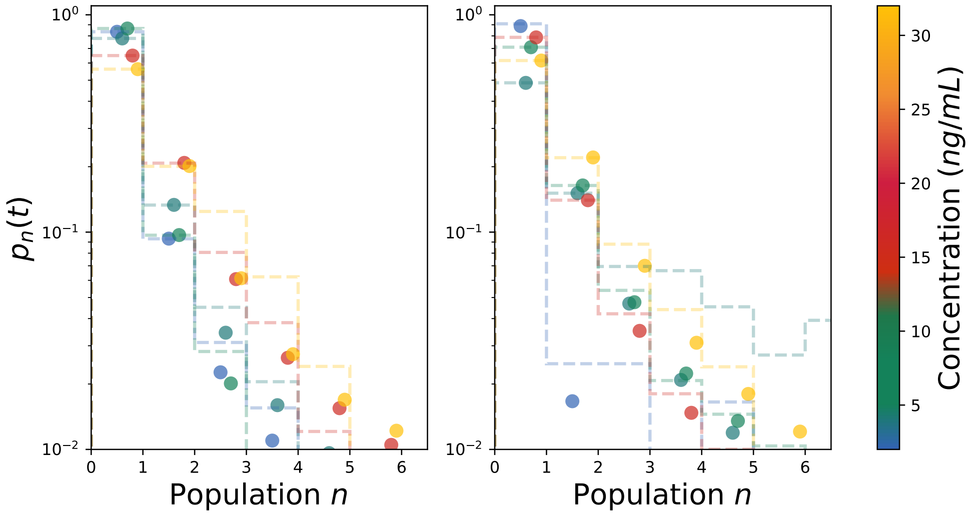


Figure 3: Comparison with 2 repetitions of the experiment. Here the parameters were computed from Eq. 39. With only one possible state we would expect a single peak at $n = 0$ and a null probability elsewhere. The data here correspond to measurements after 24 $h$, which is assumed to be a long time compared to the inverse growth or death rates.

#### 1.3 Stochastic birth process

Heredity might strongly influence phenotypes of the daughter cells, controlled by the concentrations of mRNA or proteins for instance [19, 20]. In this section we consider a different birth process shown in Fig. 4. The notations remain the same as in the previous model. We modify the model for division by assuming that each daughter cell will stay in its mother's state with probability $p$ and acquire a different state drawn from the distribution $(\pi_i)_{1\leq i\leq P}$ with probability $1-p$. This way, we can vary the strength of heredity by varying $p$. Here heredity concerns the correlations between mother's and daughter's bacterial states rather than the correlation between mother's and daughter's division sizes or interdivision times [12]. We treat each daughter cells as independent. Therefore, the new master equation is

$$\begin{aligned} p(\boldsymbol{n}, t+dt, c) - p(\boldsymbol{n}, t, c) = &\sum_i \nu_i \left[(n_i+1)p(\boldsymbol{n}+\boldsymbol{u_i}, t, c) - n_i p(\boldsymbol{n}, t, c)\right] dt \\ &+ \sum_{i,j,k} (1-p)^2 \pi_j \pi_k \lambda_i \left[(n_i + 1 - \delta_{i,j} - \delta_{i,k}) p(\boldsymbol{n} - \boldsymbol{u_i} - \boldsymbol{u_j} + \boldsymbol{u_k}, t, c) - n_i p(\boldsymbol{n}, t, c)\right] dt \\ &+ \sum_{i,j} 2p(1-p)\pi_j \lambda_i \left[(n_i - \delta_{i,j}) p(\boldsymbol{n} - \boldsymbol{u_j}, t, c) - n_i p(\boldsymbol{n}, t, c)\right] dt \\ &+ \sum_i p^2 \lambda_i \left[(n_i - 1)\, p(\boldsymbol{n} - \boldsymbol{u_i}, t, c) - n_i p(\boldsymbol{n}, t, c)\right] dt. \end{aligned} \tag{40}$$

Using, the generating function as above, we find that

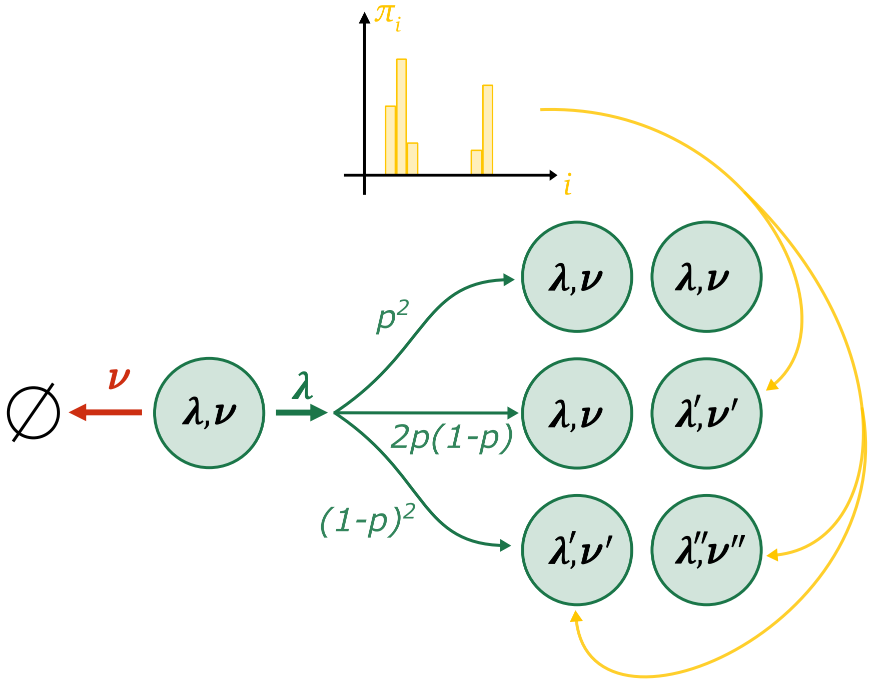


Figure 4: Death/birth model with stochastic jumps between bacterial states, drawn from a distribution at each division with a probability $p$.

$$
\begin{aligned}
\frac{\partial F(\boldsymbol{z},t,c)}{\partial t} &= \sum_i \nu_i(1-z_i)\partial_{z_i}F + \sum_{i,j,k}(1-p)^2\pi_j\pi_k\lambda_i(z_jz_k - z_i)\partial_{z_i}F \\
&+ \sum_{i,j} 2p(1-p)\pi_j\lambda_i z_i(z_j-1)\partial_{z_i}F + \sum_i p^2\lambda_i z_i(z_i-1)\partial_{z_i}F \\
&= \sum_i \left( \left(\nu_i - p^2\lambda_i z_i\right)(1-z_i) + (1-p)^2\left(\sum_{i,j,k}\pi_j\pi_k\lambda_i(z_jz_k - z_i)\right) + 2p(1-p)\left(\sum_j \pi_j(z_j-1)\right)\lambda_i z_i\right)\partial_{z_i}F \\
&= \sum_i -p^2\lambda_i\left( z_i^2 - z_i\left(\frac{1-p}{p}\left(\frac{1+p}{p} - 2\sum_j \pi_j z_j\right) + \frac{\nu_i}{\lambda_i p^2} + 1\right) + \frac{\nu_i}{\lambda_i p^2} + \frac{(1-p)^2}{p^2}\left(\sum_j \pi_j z_j\right)^2\right)\partial_{z_i}F.
\end{aligned}
\tag{41}
$$

Using again the method of characteristics, we look for variables $u, \{v_i\}_i$ such that

$$
\begin{aligned}
&\frac{dt}{du} = 1 \\
&\quad \forall i > 1,\ \frac{dz_1}{du} = 0 \\
&\frac{dz_1}{du} = -\lambda_1 \left( (p^2{+}2\pi_1 p(1-p){+}\pi_1^2(1-p)^2) z_1^2 {-} z_1 \left( 1{+}q_1{-}2(1-p)\sum_{j\neq 1} \pi_j z_j (p + \pi_1(1-p)) \right) {+} q_1 {+} (1-p)^2 \left( \sum_{j\neq 1} \pi_j z_j \right)^2 \right),
\end{aligned}
\tag{42}
$$

which ensures that $\frac{dF}{du} = 0$, with the initial parametric conditions

$$
\begin{aligned}
t(u = 0, \{v_i\}_i) &= 0 \\
z_i(u = 0, \{v_j\}_j) &= v_i \\
F(\{v_i\}_i, u = 0, c) &= \prod_{k=1}^{P} v_k^{n_0(k)}.
\end{aligned}
\tag{43}
$$

We define $q_i = \nu_i/\lambda_i$ and

$$
\begin{aligned}
\Delta_i &= \left( 1{+}q_i{-}2(1-p)\sum_{j\neq i} \pi_j z_j (p + \pi_i(1-p)) \right)^2 - 4\left(p^2{+}2\pi_i p(1-p){+}\pi_i^2(1-p)^2\right) \left( q_i + (1-p)^2 \left( \sum_{j\neq i} \pi_j z_j \right)^2 \right) \\
A_i &= \frac{1}{2} \left( 1{+}q_i{-}2(1-p)\sum_{j\neq i} \pi_j z_j (p + \pi_i(1-p)) \right) \\
B_i &= \frac{1}{2}\sqrt{\Delta_i}.
\end{aligned}
\tag{44}
$$

Such that

$$
A_i^2 = B_i^2 + \left(p^2{+}2\pi_i p(1-p){+}\pi_i^2(1-p)^2\right) \left( q_i + (1-p)^2 \left( \sum_{j\neq i} \pi_j z_j \right)^2 \right). \tag{45}
$$

**Separation of timescales** Assuming the same separation of timescales as in the first model,

$$
\begin{aligned}
&\forall j > 1,\ \frac{dz_j}{du} = 0 \\
&\left( \frac{1}{z_1 - \frac{A_1 + B_1}{p^2+2\pi_1 p(1-p)+\pi_1^2(1-p)^2}} - \frac{1}{z_1 - \frac{A_1 - B_1}{p^2+2\pi_1 p(1-p)+\pi_1^2(1-p)^2}} \right) dz_1 = -2\lambda_1 B_1 du,
\end{aligned}
\tag{46}
$$

then the resolution is the same as above, and we obtain

$$F(\{z_i\},t,c) = \left( \frac{z_1\left(\coth(\lambda_1 B_1 t) - \frac{A_1}{B_1}\right) - \frac{B_1}{p^2+2\pi_1 p(1-p)+\pi_1^2(1-p)^2}\left(1-\frac{A_1^2}{B_1^2}\right)}{\frac{A_1}{B_1} + \coth(\lambda_1 B_1 t) - (p^2 + 2\pi_1 p(1-p) + \pi_1^2(1-p)^2)\frac{z_1}{B_1}} \right) \prod_{j>1} z_j^{n_0(j)}. \quad (47)$$

In the limit of strong heredity $p \to 1$, this simplifies to

$$F(\{z_i\},t,c) = \left( \frac{z_1\left(\coth(\lambda_1 B_1 t) - \frac{A_1}{B_1}\right) - \frac{B_1}{(1-2(1-p)(1-\pi_1))}\left(1-\frac{A_1^2}{B_1^2}\right)}{\frac{A_1}{B_1} + \coth(\lambda_1 B_1 t) - (1-2(1-p)(1-\pi_1))\frac{z_1}{B_1}} \right) \prod_{j>1} z_j^{n_0(j)}, \quad (48)$$

with

$$\begin{aligned} A_1 &= \frac{1}{2}\left(1 + q_1 - 2(1-p)\sum_{j\neq 1}\pi_j z_j\right) \\ B_1 &= \frac{1}{2}\sqrt{(1-q_1)^2 + 4(1-p)\left(2q_1(1-\pi_1) - (1+q_1)\sum_{j\neq 1}\pi_j z_j\right)}. \end{aligned} \quad (49)$$

##### 1.3.1 Two states

We assume a bimodal distribution of bacterial states as earlier. We can compute the evolution of the distribution for different values of $p$. In Figs. 5 and 6. We observe that $p$ strongly modifies the evolutions of the distributions, but does not modify the survival probability. It mostly affects the time to reach a steady state as increasing $p$ seems to increase the timescale over which the distribution stabilizes.

##### 1.3.2 Starting from a Poisson binomial distribution

Now we assume that the initial distribution is a Poisson binomial with a parameters $(\mu, \pi_1)$, we typically obtain Figs. 7 and 8 respectively for small $p$ (weak heredity) and large $p$ (strong heredity). We can write the extinction probability in this case

$$p_0(t,c) = \sum_{n_0(1),n_0(2)=0,0}^{\infty} p_{n_0(1),n_0(2)} \left(\frac{2q_1}{1+q_1+2B_1\coth(\lambda_1 B_1 t)}\right)^{n_0(1)} \delta_{n_0(2),0}, \quad (50)$$

which yields

$$\begin{aligned} p_0(0,c) &= e^{-\mu} \\ p_0(t,c) &= \exp\left(\mu\left(\frac{2\pi_1 q_1}{1+q_1+2B_1\coth(\lambda_1 B_1 t)} - 1\right)\right), \end{aligned} \quad (51)$$

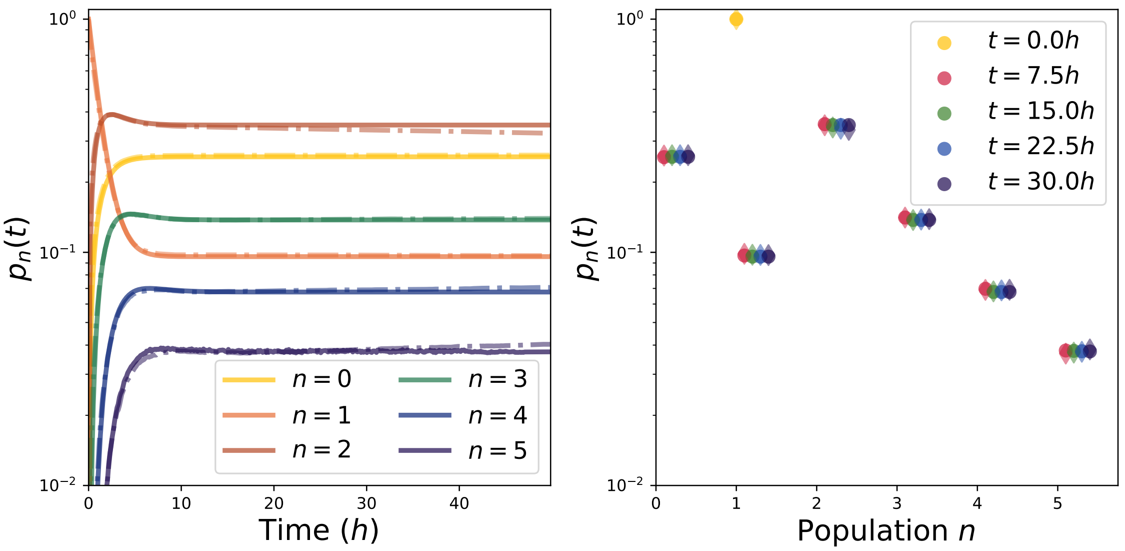


Figure 5: Death/birth model with stochastic jumps between bacterial states, drawn from a distribution at each division with a probability $p = 0.1$ and $\pi_1 = 0.3$.

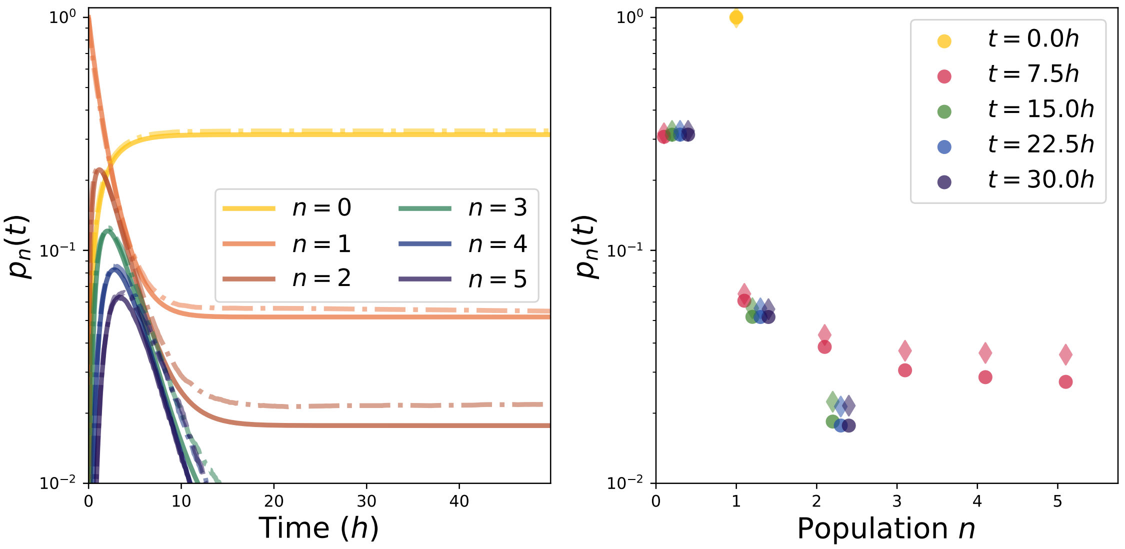


Figure 6: Death/birth model with stochastic jumps between bacterial states, drawn from a distribution at each division with a probability $p = 0.9$ and $\pi_1 = 0.3$.

**Effect of heredity and bimodality on extinction** We can study the influence of the parameters $p$ and $\pi_1$ on the extinction probability. To do so, we study $-\ln(p_0(t))/\mu$, where heredity has a weak effect on extinction probability. In particular, for $p \sim 1$ and $\pi_1 \to 0$, the population mostly contains individuals of type 2 thus reducing the extinction probability.

At large times $t \to \infty$

$$p_0(\infty, c) = \exp\left(\mu\left(\frac{2\pi_1 q_1}{1 + q_1 + 2B_1} - 1\right)\right), \tag{52}$$

so that, defining

$$\begin{aligned} r_{00} &= \frac{\ln(p_0(\infty, c))}{\ln(p_0(0, c))} \\ &= 1 - \frac{2\pi_1 q_1}{1 + q_1 + \sqrt{(1+q_1)^2 - 4q_1(p^2 + 2\pi_1 p(1-p) + \pi_1^2(1-p)^2)}}, \end{aligned} \tag{53}$$

we determine a first relationship

$$q_1 = \frac{(1 - r_{00})^2}{\pi_1(1 - r_{00} - \pi_1)}\left(p^2 + 2\pi_1 p(1-p) + \pi_1^2(1-p)^2 - \frac{\pi_1}{1 - r_{00}}\right), \tag{54}$$

which, in the strong heredity limit $p \to 1$, tends to

$$q_1 = \frac{(1 - r_{00})^2}{\pi_1(1 - r_{00} - \pi_1)}\left(1 - 2(1-\pi_1)(1-p) - \frac{\pi_1}{1 - r_{00}}\right), \tag{55}$$

**Effect of heredity and bimodality on the the ratio $p_1/p_0$** In addition we can compute the dynamics of $p_1(t, c)$ under separation of timescales,

$$\begin{aligned} \frac{p_1(t,c)}{p_0(t,c)} =& \mu\left(\pi_2 + \pi_1\left[\frac{\coth\left(\lambda_1 B_1 t\right)^2 - 1}{\left(\frac{A_1}{B_1} + \coth\left(\lambda_1 B_1 t\right)\right)^2} - \frac{\partial B_1}{\partial z_2}\frac{\frac{q_1}{B_1^2}\left(\frac{A_1}{B_1} + \coth\left(\frac{\lambda_1 B_1 t}{2}\right)\right) + \frac{\lambda_1 B_1 t\left(\frac{A_1^2}{B_1^2} - 1\right)\left(1 - \coth(\lambda_1 B_1 t)^2\right)}{p^2 + 2\pi_1 p(1-p) + \pi_1^2(1-p)^2}}{\left(\frac{A_1}{B_1} + \coth\left(\lambda_1 B_1 t\right)\right)^2}\right]\right) \\ =& \mu\left(\pi_2 + \pi_1\left[\frac{\coth\left(\lambda_1 B_1 t\right)^2 - 1}{\left(\frac{A_1}{B_1}\coth\left(\lambda_1 B_1 t\right)\right)^2}\right.\right. \\ & \left.\left. + \pi_2(1+q_1)(1-p)(2p + \pi_1(1-p))\frac{\frac{q_1}{B_1^2}\left(\frac{A_1}{B_1} + \coth\left(\lambda_1 B_1 t\right)\right) - \frac{\lambda_1 B_1 t\left(\frac{A_1^2}{B_1^2} - 1\right)\left(\coth(\lambda_1 B_1 t)^2 - 1\right)}{p^2 + 2\pi_1 p(1-p) + \pi_1^2(1-p)^2}}{4B_1\left(\frac{A_1}{B_1} + \coth\left(\lambda_1 B_1 t\right)\right)^2}\right]\right). \end{aligned} \tag{56}$$

From this formula, we define

$$\begin{aligned} r_{01} &= \frac{p_1(\infty, c)}{\mu p_0(\infty, c)} \\ &= (1 - \pi_1)\left(1 + \frac{\pi_1 q_1 (1+q_1)(1-p)(2p + \pi_1(1-p))}{4B_1^2\left(A_1 + B_1\right)}\right). \end{aligned} \tag{57}$$

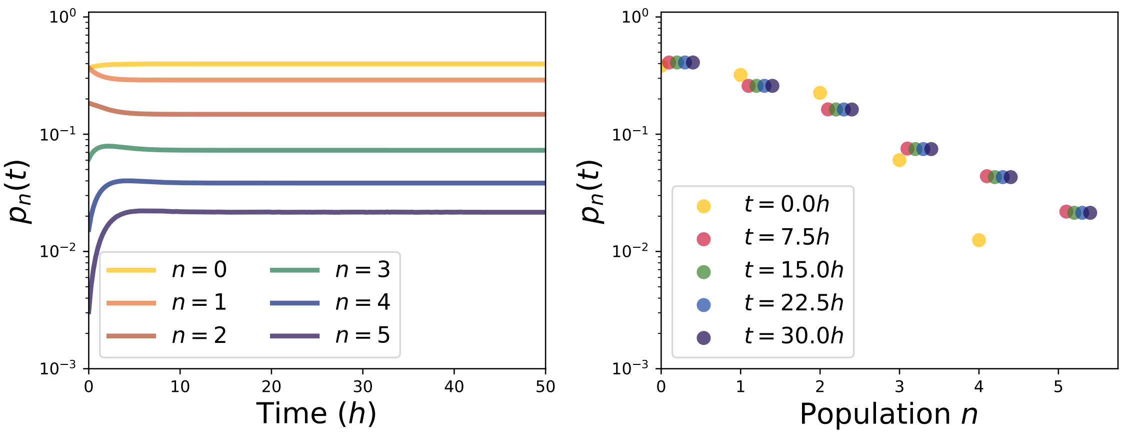


Figure 7: Death/birth model with stochastic jumps between bacterial states, drawn from a distribution at each division with a probability $p = 0.1$, starting from a Poisson binomial distribution with parameters $\mu = 1, \pi_1 = 1 - \pi_2 = 0.3$.

For strong heredity $p \to 1$, we find

$$r_{01} = (1-\pi_1)\left(1 + \frac{4\pi_1 q_1(1+q_1)(1-p)}{(1-q_1)^2\,(1+q_1+|1-q_1|)}\right). \tag{58}$$

#### 2 General correlated birth process

In this section we consider a different birth process shown in Fig. 9. We modify the equation describing birth by assuming that both daughter cells of a mother in state $i$ will acquire state $j$ with probability $\pi_{ij}$. Therefore, the new master equation is

$$\begin{aligned} p(\boldsymbol{n}, t+dt, c) - p(\boldsymbol{n}, t, c) = &\sum_i \nu_i \left[(n_i+1)p(\boldsymbol{n}+\boldsymbol{u_i}, t, c) - n_i p(\boldsymbol{n}, t, c)\right] dt \\ &+ \sum_{i,j,k} \pi_{i,j}\pi_{i,k}\lambda_i \left[(n_i+1-\delta_{i,j}-\delta_{i,k})p(\boldsymbol{n}-\boldsymbol{u_i}-\boldsymbol{u_j}+\boldsymbol{u_k}, t, c) - n_i p(\boldsymbol{n}, t, c)\right] dt. \end{aligned} \tag{59}$$

Using, the generating function as above, we find that

$$\begin{aligned} \frac{\partial F(\boldsymbol{z}, t, c)}{\partial t} &= \sum_i \nu_i(1-z_i)\partial_{z_i}F + \sum_{i,j,k}\pi_{i,j}\pi_{i,k}\lambda_i(z_j z_k - z_i)\partial_{z_i}F \\ &= \sum_i \left(\nu_i(1-z_i) + \sum_{i,j,k}\pi_{i,j}\pi_{i,k}\lambda_i(z_j z_k - z_i)\right)\partial_{z_i}F \end{aligned} \tag{60}$$

Using again the method of characteristics, we look for variables $u, \{v_i\}_i$ such that

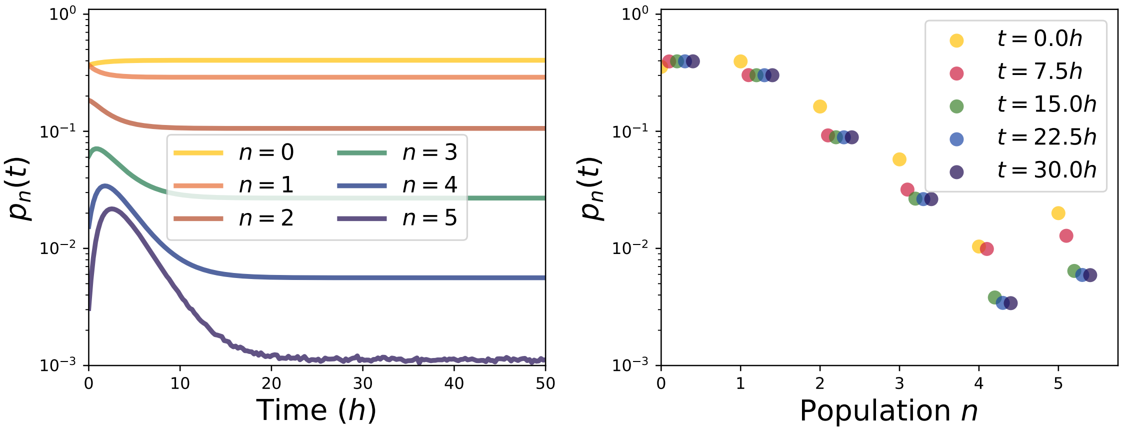


Figure 8: Death/birth model with stochastic jumps between bacterial states, drawn from a distribution at each division with a probability $p = 0.9$, starting from a Poisson binomial distribution with parameters $\mu = 1, \pi_1 = 1 - \pi_2 = 0.3$.

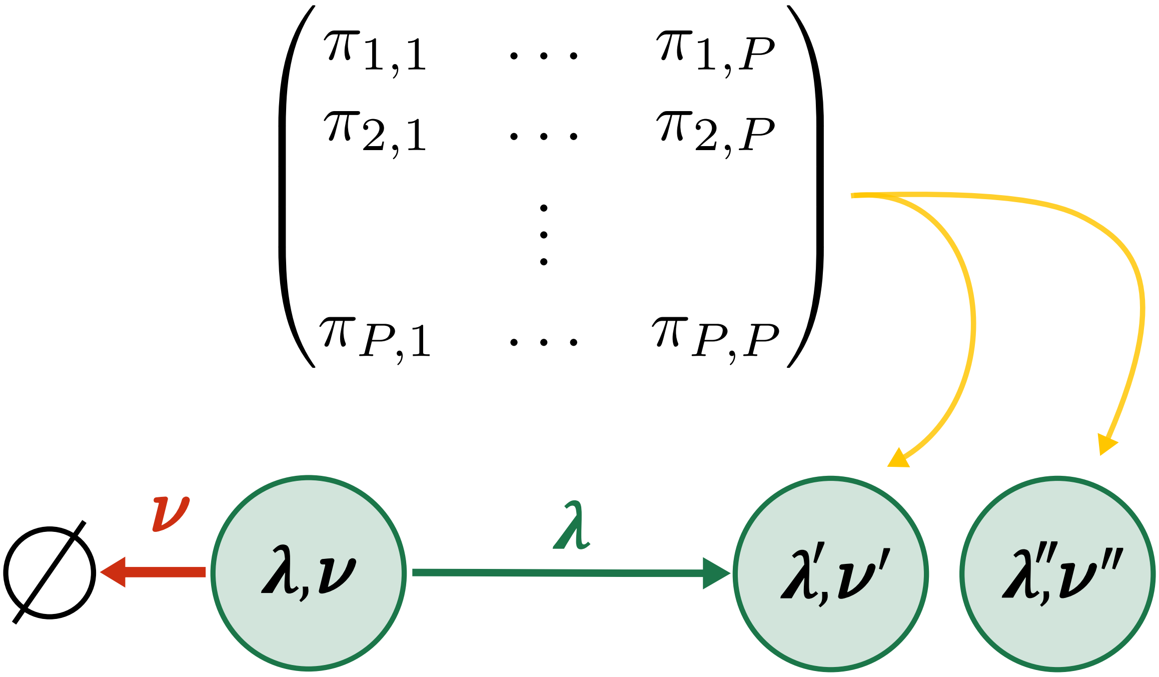


Figure 9: Death/birth model with a heredity matrix $\pi_{i,j}$. The sums over the lines $\forall i$, $\sum_j \pi_{ij} = 1$.

$$
\begin{aligned}
&\frac{dt}{du} = 1 \\
\forall i,\ &\frac{dz_i}{du} = -\lambda_i \left( \pi_{i,i}^2 z_i^2 - z_i \left( 1 + q_i - 2\sum_{j\neq i} \pi_{i,j}\pi_{i,i} z_j \right) + q_i + \left( \sum_{j\neq i} \pi_{i,j} z_j \right)^2 \right),
\end{aligned}
\tag{61}
$$

which ensures that $\frac{dF}{du} = 0$, with the initial parametric conditions

$$
\begin{aligned}
t(u = 0, \{v_i\}_i) &= 0 \\
z_i(u = 0, \{v_j\}_j) &= v_i \\
F(\{v_i\}_i, u = 0, c) &= \prod_{k=1}^{P} v_k^{n_0(k)}.
\end{aligned}
\tag{62}
$$

We define $q_i = \nu_i/\lambda_i$ and

$$
\begin{aligned}
\Delta_i &= \left( 1 + q_i - 2\sum_{j\neq i} \pi_{i,i}\pi_{i,j} z_j \right)^2 - 4\pi_{i,i}^2 \left( q_i + \left( \sum_{j\neq i} \pi_{i,j} z_j \right)^2 \right) \\
A_i &= \frac{1}{2} \left( 1 + q_i - 2\sum_{j\neq i} \pi_{i,i}\pi_{i,j} z_j \right) \\
B_i &= \frac{1}{2}\sqrt{\Delta_i},
\end{aligned}
\tag{63}
$$

such that

$$
A_i^2 = B_i^2 + \pi_{i,i}^2 \left( q_i + \left( \sum_{j\neq i} \pi_{i,j} z_j \right)^2 \right). \tag{64}
$$

Using separation of timescales

$$
\begin{aligned}
&\forall j > 1,\ \frac{dz_j}{du} = 0 \\
&\left( \frac{1}{z_1 - \frac{A_1+B_1}{\pi_{1,1}^2}} - \frac{1}{z_1 - \frac{A_1-B_1}{\pi_{1,1}^2}} \right) dz_1 = -2\lambda_1 B_1 du,
\end{aligned}
\tag{65}
$$

then the resolution is the same as above, and we obtain

$$
F(\{z_i\}, t, c) = \left( \frac{z_1 \left( \coth(\lambda_1 B_1 t) - \frac{A_1}{B_1} \right) - \frac{B_1}{\pi_{1,1}^2} \left( 1 - \frac{A_1^2}{B_1^2} \right)}{\frac{A_1}{B_1} + \coth(\lambda_1 B_1 t) - \pi_{1,1}^2 \frac{z_1}{B_1}} \right)^{n_0(1)} \prod_i z_i^{n_0(i)}, \tag{66}
$$

which behaves as in the previous model and heredity is set by the matrix $\pi_{i,j}$ (the larger the diagonal elements the stronger the heredity) instead of the parameter $p$. The trace of $\pi$ thus quantifies heredity.

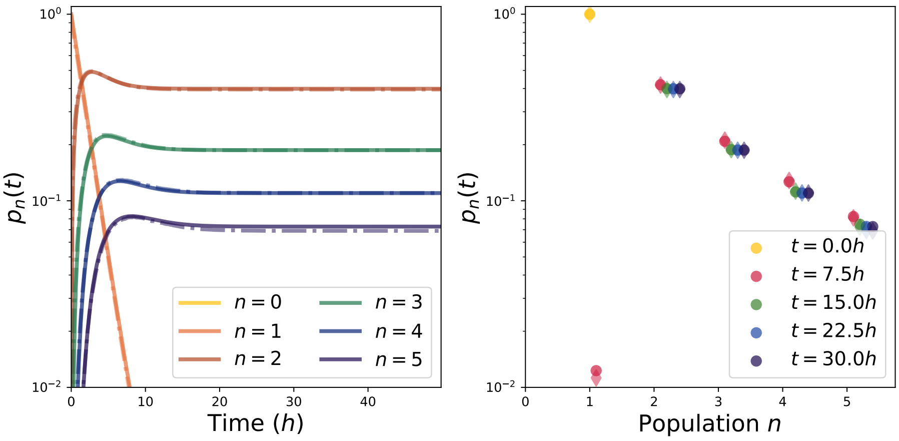


Figure 10: Death/birth model with stochastic jumps between bacterial states drawn from a distribution at each division with a probability $p = 0.1$ and $\pi_1 = 0.3$, where the dormant state represent dead cells (still counted here).

### 3 Case of failure to identify dead cells

**Birth/Death model with one daughter similar to the mother** Now if we assume that dead cells are still counted together with living cells, we would only observe transitions from the active state 1 to itself or to the dormant/dead state 2, and $\nu_1 = \nu_2 = 0$, and $\lambda_2 = 0$. Then in such a setting, since active cells always give at least one active daughter, the population of a droplet containing at least one active cell cannot reach extinction. This model is therefore not adapted to describe transitions to a dead state.

**Birth/death model with partial heredity** With the second model (where heredity is controlled by a parameter $p$), we have $\nu_1 = \nu_2 = 0$, and $\lambda_2 = 0$. In this case, we obtain the same results with $A_1 = (1 - 2(1-p)\pi_2 z_2)/2$ and $B_1 = \sqrt{1 - 4(1-p)\pi_2 z_2}/2$ (in particular $A_1 = B_1$ in the limit of strong heredity $p \to 1$). We show the time dependent probabilities in Fig.10. The initial condition is deterministic here (one cell in the active state), thus at least one division occurs (and $p_1(t)$ goes to 0). A plateau is reached after a few hours, so that the probability to end up with a few individuals at large time is non zero here as well. Heredity plays the same role, and a strong heredity ensures that most cells remain in an active state.

The observed dynamics depends on the initial condition, which is that the droplet contains one active cell. Due to this initial condition, there will be at least one division observed, explaining why the peak at 1 is decreasing with time and no peak is observed at 0 (extinction, in the sense that no individual is counted, is not possible in this case). The probabilities to observe larger populations reach a steady state corresponding to droplets where all individuals are in the dormant/dead state (steady absorbing state). We would recover a tailed distribution by starting with a stochastic initial condition, where the peak for extinct populations (probability of having an empty droplet) is constant over time.

**General correlated model with a correlation matrix** Still with the same assumption and relying on the transition matrix $\pi_{ij}$, we would only see transitions from the active state 1 to itself or to the dormant/dead state 2, meaning that $\pi_{21} = 0$, and $\nu_1 = \nu_2 = 0$. In this case we obtain the same results with $A_1 = (1 - 2\pi_{11}\pi_{12}z_2)/2$ and $B_1 = \sqrt{1 - 4\pi_{11}\pi_{12}z_2}/2$ (here as well, $A_1 = B_1$ in the limit of strong heredity $p \to 1$).

### 4 Transition dependence on drug concentration

In principle, the transition between dormant/inhibited and active states should depend on antibiotic concentration. In particular, we expect that transition towards dormant/inhibited increases with drug concentration. Growth rates also depend on antibiotic concentration [21, 22], and bistability should appear in the irreversible limit where antibiotics accumulate inside the cell. The simplest model we can imagine for the dependency of $\pi_1, \pi_2$ on drug concentration $c$ (for small $c$) is a linear by parts model

$$\pi_2 = \begin{cases} 0 & \text{for } c < c_0 \\ \frac{c-c_0}{c_1-c_0} & \text{for } c_0 \leq c < c_1 \\ 1 & \text{for } c \geq c_1 \end{cases}, \tag{67}$$

where $c_0 < c_1$, $c_0$ is the first bifurcation point and $c_1$ the second bifurcation point. In the irreversible limit, the growth rate as a function of antibiotic concentration has three available values [21]

$$\frac{\lambda}{\lambda_0} \in \left\{0, \frac{1}{2}\left(1 - \sqrt{1 - \frac{c}{IC_{50}}}\right), \frac{1}{2}\left(1 + \sqrt{1 - \frac{c}{IC_{50}}}\right)\right\}, \tag{68}$$

where $IC_{50}$ is the half inhibitory concentration [21, 22] that is the concentration of antibiotics for which the growth rate is half the drug free growth rate. Only 0 and $\left(1 + \sqrt{1 - c/IC_{50}}\right)/2$ are stable, and in this case we find $c_0 \sim 0$ and $c_1 \sim IC_{50}$, meaning that

$$\pi_2 = \begin{cases} \frac{c}{IC_{50}} & \text{for } c < IC_{50} \\ 1 & \text{for } c \geq IC_{50} \end{cases}. \tag{69}$$

We perform a linear regression on these data (using the scipy function linregress). In practice, the linear regression can be accepted with p-values of 0.03 for the first dataset and 0.07 for the second dataset for small $c \ll IC_{50}$, as shown on Fig.11 (using Wald test with t-distributions where the null hypothesis is that the slope is 0, that is testing the t-ratio $\hat{p}/\sigma_{\hat{p}}$ where $\hat{p}$ is the gradient, serving as an estimator of the slope, which should follow a t-distribution with $n_{\text{sample}} - 2$ degrees of freedom, due to the $n_{\text{sample}} - 2$ estimates of the gradient). We conclude that there is a positive correlation between the parameter $\pi_2$ and the antibiotic concentration.

On the contrary, $q_1$ is not well described by a linear model, and there is no clear trend appearing for $q_1$ as a function of the drug concentration. This complex behavior comes from the fact that both the death and division rates depend on the antibiotic concentration.

### References


[1] L. Le Quellec, A. Aristov, S. Gutiérrez Ramos, G. Amselem, J. Bos, Z. Baharoglu, D. Mazel, and C. N. Baroud. Measuring single-cell susceptibility to antibiotics within monoclonal bacterial populations. *PLoS One*, 19(8):e0303630, 2024.

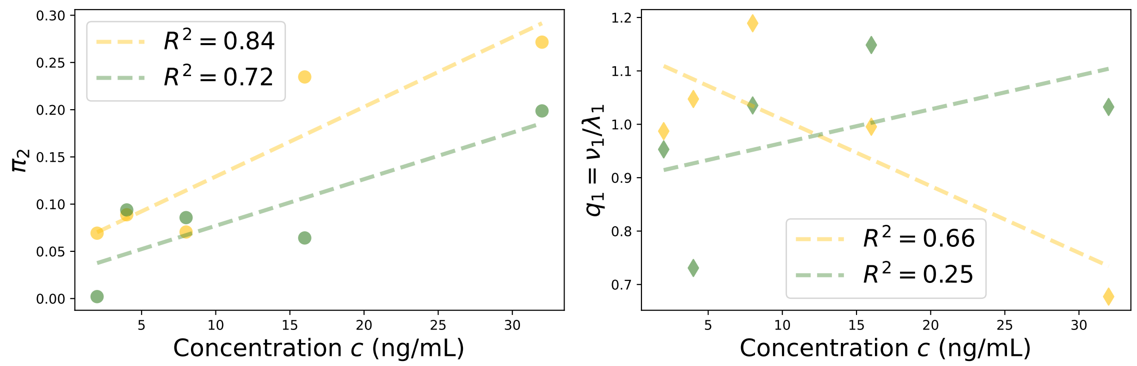


Figure 11: Parameters as functions of antibiotic concentration


[2] A. Barizien. *Studying the variability of bacterial growth in microfluidic droplets.* Theses, Univ. Paris Saclay, 2019.

[3] E. Maikranz, A. Aristov, L. Le Quellec, and C. Baroud. Extending digital biology: bacterial survival and morphological heterogeneity under antibiotic stress, 2025.

[4] G. Amselem, C. Guermonprez, B. Drogue, S. Michelin, and C. N. Baroud. Universal microfluidic platform for bioassays in anchored droplets. *Lab Chip*, 16(21):4200–4211, 2016.

[5] A. Barizien, M. S. Suryateja Jammalamadaka, G. Amselem, and C. N. Baroud. Growing from a few cells: combined effects of initial stochasticity and cell-to-cell variability. *J. R. Soc. Interface*, 16(153):20180935, 2019.

[6] A. Roy and S. Klumpp. Simulating Genetic Circuits in Bacterial Populations with Growth Heterogeneity. *Biophys. J.*, 114(2):484–492, 2018.

[7] D. Huh and J. Paulsson. Non-genetic heterogeneity from stochastic partitioning at cell division. *Nat. Genet.*, 43:95–100, 2011.

[8] D. Huh and J. Paulsson. Random partitioning of molecules at cell division. *Proc. Natl. Acad. Sci. U.S.A.*, 108:15004–15009, 2011.

[9] Q. Chai, B. Singh, K. Peisker, N. Metzendorf, X. Ge, S. Dasgupta, and S. Sanyal. Organization of Ribosomes and Nucleoids in Escherichia coli Cells during Growth and in Quiescence *. *J. Biol. Chem.*, 289:11342–11352, 2014.

[10] A. Papagiannakis, Q. Yu, S. K. Govers, W. Lin, N. S. Wingreen, and C. Jacobs-Wagner. Nonequilibrium polysome dynamics promote chromosome segregation and its coupling to cell growth in Escherichia coli. *eLife*, 14:RP104276, 2025.

[11] A. M. Miangolarra, S. H. Li, J. Joanny, N. S. Wingreen, and M. Castellana. Steric interactions and out-of-equilibrium processes control the internal organization of bacteria. *Proc. Natl. Acad. Sci. U.S.A.*, 118(43):e2106014118, 2021.

[12] J. Lin and A. Amir. From single-cell variability to population growth. *Phys. Rev. E*, 101:012401, 2020.

[13] J. Grilli, C. Cadart, G. Micali, M. Osella, and M. Cosentino Lagomarsino. The Empirical Fluctuation Pattern of E. coli Division Control. *Front. Microbiol.*, Volume 9 - 2018, 2018.

[14] M. Arnoldini, I. A. Vizcarra, R. Peña-Miller, N. Stocker, M. Diard, V. Vogel, R. E. Beardmore, W.-D. Hardt, and M. Ackermann. Bistable Expression of Virulence Genes in Salmonella Leads to the Formation of an Antibiotic-Tolerant Subpopulation. *PLoS Biol.*, 12(8):e1001928, 2014.

[15] J. Karslake, J. Maltas, P. Brumm, and K. B. Wood. Population Density Modulates Drug Inhibition and Gives Rise to Potential Bistability of Treatment Outcomes for Bacterial Infections. *PLoS Comput. Biol.*, 12:e1005098, 2016.

[16] P. Patra and S. Klumpp. Population Dynamics of Bacterial Persistence. *PLoS One*, 8(5):e62814, 2013.

[17] T. Dörr, M. Vulić, and K. Lewis. Ciprofloxacin causes persister formation by inducing the TisB toxin in Escherichia coli. *PLoS Biol.*, 8(2):e1000317, 2010.

[18] T. Dörr, K. Lewis, and M. Vulić. SOS response induces persistence to fluoroquinolones in Escherichia coli. *PLoS Genet.*, 5(12):e1000760, 2009.

[19] D. Antunes and A. Singh. Quantifying gene expression variability arising from randomness in cell division times. *J. Math. Biol.*, 71(2):437–463, 2015.

[20] B. Cerulus, A. M. New, K. Pougach, and K. J. Verstrepen. Noise and Epigenetic Inheritance of Single-Cell Division Times Influence Population Fitness. *Curr. Biol.*, 26(9):1138–1147, 2016.

[21] P. Greulich, M. Scott, M. R. Evans, and R. J. Allen. Growth-dependent bacterial susceptibility to ribosome-targeting antibiotics. *Mol. Syst. Biol.*, 11(3):796, 2015.

[22] B. Ledoux and D. Lacoste. Inhibition of bacterial growth by antibiotics: a minimal model. *Phys. Biol.*, 22(6):066007, 2025.
